# Disrupted choline clearance and sustained acetylcholine release *in vivo* by a common choline transporter coding variant associated with poor attentional control in humans

**DOI:** 10.1101/2021.06.27.450076

**Authors:** Eryn Donovan, Cassandra Avila, Sarah Klausner, Vinay Parikh, Cristina Fenollar-Ferrer, Randy D. Blakely, Martin Sarter

## Abstract

Transport of choline via the neuronal high-affinity choline transporter (CHT; *SLC5A7)* is essential for cholinergic terminals to synthesize and release acetylcholine (ACh). In humans, we previously demonstrated an association between a common CHT coding substitution (rs1013940; Ile89Val) and reduced attentional control as well as attenuated frontal cortex activation. Here, we used a CRISPR/Cas9 approach to generate mice expressing the I89V substitution and assessed, *in vivo,* CHT-mediated choline transport, and ACh release. Relative to wild type (WT) mice, CHT-mediated clearance of choline in mice expressing one or two Val89 alleles was reduced by over 80% cortex and by over 50% in striatum. Choline clearance in CHT Val89 mice was further reduced by neuronal inactivation. Deficits in ACh release, 5 and 10 min after repeated depolarization at a low, behaviorally relevant frequency, support an attenuated reloading capacity of cholinergic neurons in mutant mice. The density of CHTs in total synaptosomal lysates and neuronal plasma-membrane-enriched fractions was not impacted by the Val89 variant, indicating a selective impact on CHT function. When challenged with a visual disruptor to reveal attentional control mechanisms, Val89 mice failed to adopt a more conservative response bias. Structural modeling revealed that Val89 may attenuate choline transport by altering conformational changes of CHT that support normal transport rates. Our findings support the view that diminished, sustained cholinergic signaling capacity underlies perturbed attentional performance in individuals expressing CHT Val89. The CHT Val89 mouse serves as a valuable model to study heritable risk for cognitive disorders arising from cholinergic dysfunction.

**Significance Statement:** Acetylcholine (ACh) signaling depends on the functional capacity of the neuronal choline transporter (CHT). Previous research demonstrated that humans expressing the common CHT coding variant Val89 exhibit attentional vulnerabilities and attenuated fronto-cortical activation during attention. Here, we find that mice engineered to express the Val89 variant exhibit reduced CHT-mediated choline clearance and a diminished capacity to sustain ACh release. Additionally, Val89 mice lack cognitive flexibility in response to an attentional challenge. These findings provide a mechanistic and cognitive framework for interpreting the attentional phenotype associated with the human Val89 variant and establish a model that permits a more invasive interrogation of CNS effects as well as the development of therapeutic strategies for those, including Val89 carriers, with presynaptic cholinergic perturbations.

## Introduction

Activation of forebrain cholinergic projection systems is necessary for a wide range of perceptual-attentional functions. In humans, disruption of ACh signaling, either by pharmacological blockade of cholinergic receptors or as a result of cholinergic cell loss, impairs the capacity to select behaviorally significant stimuli for behavioral control, to filter distractors, and to perform under dual-task conditions (e.g., Thienel et al., 2009; Mentis et al., 2001; Kim et al., 2019; Albin et al., 2018; Yarnall et al., 2011; Perry and Hodges, 1999). Results from experiments in non-human primates and rodents have confirmed the essential role of forebrain cholinergic activity for performing a broad range of attentional tasks (e.g., Muir et al., 1992; Voytko et al., 1994; McGaughy et al., 1996; Turchi and Sarter, 2000, 1997) and specified underlying cholinergic signaling characteristics and postsynaptic mechanisms (e.g., Venkatesan and Lambe, 2020; Howe et al., 2017; Gritton et al., 2016; Chen et al., 2015; Parikh et al., 2007; Sarter and Lustig, 2020; Nair et al., 2018; Laszlovszky et al., 2020; Yang et al., 2021; Lu et al., 2020).

The sodium-dependent, high-affinity, neuronal choline transporter (CHT) is essential for sustaining the capacity of presynaptic cholinergic neurons to synthesize and release ACh (Yamamura and Snyder, 1972; Guyenet et al., 1973; Simon et al., 1976; Ennis and Blakely, 2016). Early experiments assessing the effects of pharmacological blockade of the CHT with the competitive CHT antagonist hemicholinium-3 (HC-3), and more recent investigations into the impact of CHT heterozygosity in transgenic mice (Parikh et al., 2013; Paolone et al., 2013b), have demonstrated that reduced CHT capacity causes attenuated cholinergic signaling and broadly disrupts perceptual-attentional functions (for reviews see Ferguson and Blakely, 2004; Sarter and Parikh, 2005; Bazalakova and Blakely, 2006; Sarter et al., 2016).

Okuda et al. (2002) discovered a common, single nucleotide polymorphism (SNP) within human CHT coding sequences (rs1013940) that produces the Ile89Val, or I89V, substitution in transmembrane domain 3, with ~8% minor allele frequency in Caucasians (1000 Genomes Project). Importantly, expression of the Val89 variant in transfected cells demonstrated reduced choline transport (40-50% of WT CHT) without changes in transporter surface expression or substrate affinity. We previously demonstrated that humans heterozygous for the Val89 variant exhibit attention control deficits when challenged with a distractor (Berry et al., 2015; Berry et al., 2014; Sarter et al., 2016). Moreover, right frontal activation during an attention-demanding task was significantly attenuated in these individuals (see also Gorka et al., 2015). Clinical significance of the expression of the CHT Val89 variant was indicated by studies showing an association with attention-deficit hyperactivity disorder (English et al., 2009) and major depression (Hahn et al., 2008).

The current experiments aimed to determine, *in vivo,* the impact of the CHT Val89 variant on cortical and striatal choline transport and cortical ACh release when introduced into mouse CHT coding sequences using CRISPR/Cas9 methods. Using these mutant mice and their Ile89 (WT) littermates, we establish a significant reduction in the capacity of Val89-expressing mice to clear choline from the extracellular space. Furthermore, successive, depolarization-evoked ACh release events, induced at a frequency mirroring cholinergic activity in attention task-performing rodents, led to a robust attenuation of ACh release in the mutant mice. These cholinergic capacity limits by the CHT Val89 variant were not associated with altered surface expression of the transporter. Consistent with attentional control deficits seen in Val89-expressing humans, a visual disruptor revealed that WT mice, but not Val89 mice, deployed attentional control mechanisms to perform a continuous signal detection task. Structural modeling studies suggested that the Val89 substitution reduces choline transport by potentially disrupting the architecture of the CHT close to the substrate binding site. Our findings provide a neurobiological and cognitive framework for the inattentive phenotype that characterizes human Val89 carriers.

## Materials and Methods

### Subjects

Mice were generated to express a valine rather than the WT isoleucine residue at amino acid 89 in the endogenous *Slc5a7* gene that encodes the CHT. Briefly, a CRISPR/Cas9 approach (Cong et al., 2013) was implemented through the Transgenic Mouse Embryonic Stem Cell Shared Resource facility at Vanderbilt University. An RNA guide expressing construct was developed by ligation of a 193 bp protospacer into the Bbsl sites of the pX330 plasmid (http://www.addgene.org/42230/) that also affords expression of human codon optimized Cas9. The protospacer contained an 82-nucleotide arm 5’ of the codon (ATT) that encodes Ile89 in exon 3 and a 93-nucleotide arm 3’ of the site in intron 4 (HDR oligos). An 18mer donor oligonucleotide (5’CC**<u>G</u>**T**<u>G</u>**GG**<u>C</u>**TATTCTCTGAGTC3’) was generated to encode a Val89 codon along with the introduction of a synonymous codon change in the adjacent codon (GG**<u>A</u>** to GG**<u>C</u>**), designed to reduce the chance of repeated CRISPR activity on the edited sequence. Base changes also afford diagnostic restriction digestion (novel Hpy99I site) to ascertain the presence of the sequence change introduced. Cas9/gRNA plasmid and donor oligonucleotide were injected into single cell C57BL6/J mouse embryos subsequently implanted into a pseudopregnant female mouse. After weaning, tissue from tail samples of pups were genotyped by restriction digest, followed by DNA sequencing of PCR fragments spanning the mutation, cloned into pGEMTEasy (Promega). From 3 different litters, 3 animals were identified as heterozygous founders carrying the intended codon mutation without other deletions apparent. Two of these founders produced pups carrying the Val89 allele, one of which served as the founder for the line studied. These animals were subsequently crossed to wild type C57BL6/J mice for eight generations before maintaining the colony as homozygous or heterozygous breeders.

The data from a total of N=117 mice (n=38 CHT^Ile/Ile^ mice, henceforth termed WT, 18 females; n=38 heterozygous CHT^Ile/Val^ mice, henceforth termed CHT I/V,18 females; and 41 homozygous CHT^Val/Val^ mice, henceforth termed CHT V/V, 17 females) were included in the final analyses. Following arrival at the University of Michigan, the CHT Val89 colony was further crossed to C57BL/6J mice for an additional four generations. Mice were housed in a temperature and humidity-controlled environment (23°C, 45%) with a 12:12 hour light/dark cycle (lights on at 7:00 A.M.). Data collection for all experiments began during the light phase. Litters were weaned at 3 weeks and group-housed with same-sex siblings for a maximum of 6 animals per cage. Genomic DNA from tail tissue samples was used to determine mouse genotype as either WT, CHT I/V or CHT V/V by Transnetyx (Cordova, TN). Animals were given food (Envigo Teklad rodent diet) and water *ad libitum.* Mice were a minimum of 8 weeks old at the beginning of experiments. All procedures were conducted in adherence with protocols approved by the Institutional Animal Care and Use Committee (IACUC) at the University of Michigan.

#### Mice used in individual experiments

Amperometric recordings of *in vivo* choline clearance were used to quantify the impact of the Val89 variant on CHT uptake efficiency under various conditions. Initial experiments determined exogenous choline clearance in cortex of WT (n=4, 3 females), CHT I/V (n=4, 1 female), and CHT V/V (n=5, 2 females) mice, and in striatum of WT striatum of WT (n=4, 2 females), CHT I/V (n=3, 2 females), and CHT V/V (n=3, 2 females) mice. A total of 15 additional mice (6 females) were used to determine choline clearance in the presence of the CHT competitive inhibitor HC-3 (n=5 WT, n=4 CHT I/V, n=6 CHT V/V; 2 females for each genotype). In separate mice (n=6, 2 per genotype, 1 female per genotype), the contribution of diffusion rate to overall choline clearance was quantified in the presence of lidocaine HCl to arrest transporter activity and, more broadly, neuronal firing (Tehovnik and Sommer, 1997; Malpeli and Schiller, 1979; Chu and Lee, 1994). To illustrate the efficacy of lidocaine, we also determined endogenous ACh release in the presence of lidocaine in one WT male mouse. To demonstrate attenuated ACh release following repeated potassium-evoked depolarizations, a total of 15 mice were used (WT, n=4, 2 females; CHT I/V, n=6, 3 females; CHT V/V, n=5, 3 females). A group of 30 mice (equal group sizes and sex distribution across genotypes) were used to assess CHT densities in total synaptosomal lysates and synaptosomal plasma membrane-enriched fractions. Data from 27 mice (WT, n=8, 3 females; CHT I/V, n=9, 4 females; CHT V/V, n=10, 2 females) that reached the acquisition criterion in a continuous signal detection task were used to characterize their attentional phenotype.

### Amperometric recordings of choline currents

The experiments described herein measured choline concentrations in the extracellular space. We determined the capacity of the CHT by assessing the effects of pressure ejections of choline on CHT-mediated choline clearance. The capacity of cholinergic neurons to release ACh was determined by assessing the effects of repeated KCl-induced depolarization on extracellular choline concentrations. Prior experiments validated the use of choline-sensitive microelectrodes to monitor ACh release in real-time, in part by demonstrating that, *in vivo,* endogenous acetylcholinesterase (AChE) completely hydrolyzes new released ACh (Parikh et al., 2013; Giuliano et al., 2008; Parikh et al., 2007; Parikh and Sarter, 2006; Parikh et al., 2004).

Choline concentrations were measured using an amperometric measurement scheme, illustrated in Figure 1. In this approach, choline oxidase (CO) is immobilized onto the surface of an electrode to oxidize choline, resulting in a byproduct of hydrogen peroxide. Electrochemical oxidation of hydrogen peroxide then results in the release of two electrons, which generate the current that is translated into the concentration of choline at the recording site based on prior *in vitro* calibrations (Burmeister et al., 2003; Burmeister et al., 2000).

**Figure 1.**
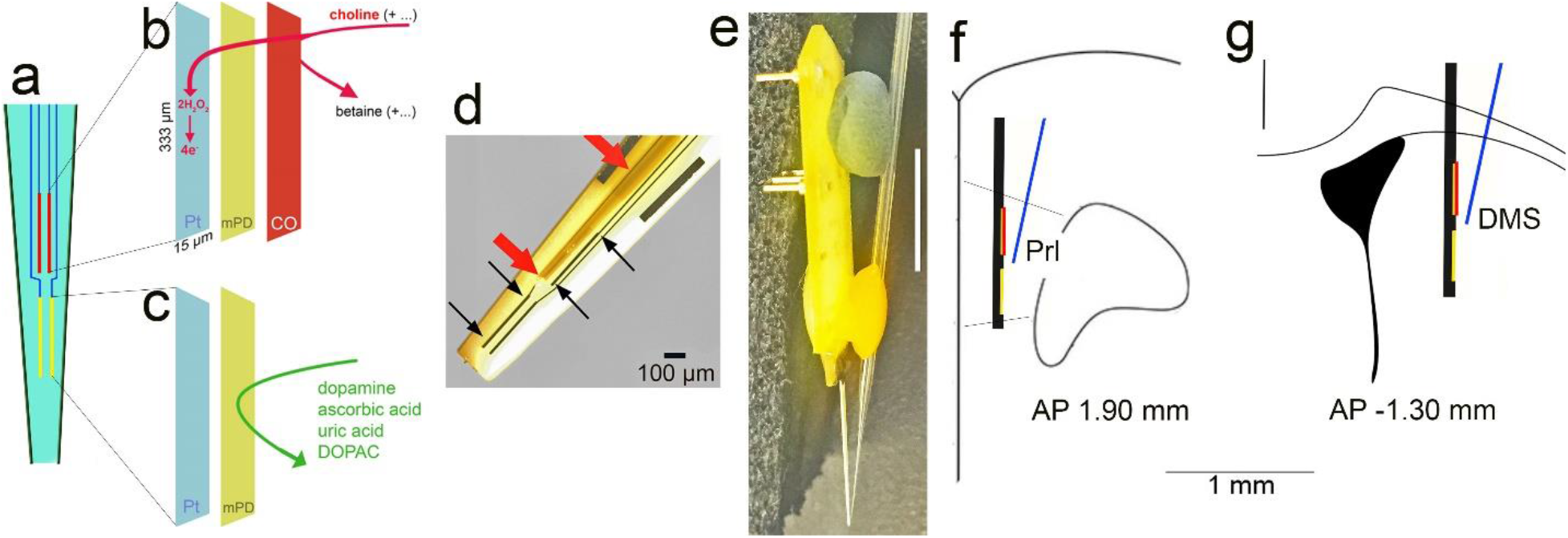
Diagram of the choline measurement scheme, the glass capillary attached to the electrode and for pressure ejections, and placement of the assembly in the medial frontal cortex. **a** shows the ceramic backbone with the 4 Platinum (Pt/Ir) recording sites, each 333 μm long and 15 μm wide, organized in two pairs (see text for references) The upper pair was used to measure choline currents (red; see also **b**) while the lower pairs served as sentinels (yellow; see also **c**). The choline-sensing scheme is illustrated in abbreviated form in b and c. Choline is oxidized by choline oxidase (CO) that was immobilized onto the upper pair of recording sites. The resulting current, over background activity, was converted to concentration (μM). Prior to immobilization of CO, the non-conducting polymer m-(1,3)-phenylenediamine (mPD) was electroplated onto the recording sites to block the transfer of small electroactive organic molecules to the Pt site. Importantly, such mPD films do not block small species such as H_2_O_2_ from reaching the Pt surface (Mitchell, 2004; Huang et al., 1993). The lower pair of recording sites (c) were coated with mPD but not CO and thus served to record the concentration of electroactive interferents, which, despite the mPD barrier, still reached the Pt sites. A top view of the electrode in d shows the Pt recording sites (black arrows), with the upper left Pt site situated underneath a glass capillary (red arrows) used to pressure eject compounds into the recording area (100 μm scale inserted). The entire electrode/glass capillary assembly is shown from the side in **e**. Note that the Pt recording sites are not visible because of low resolution (1-cm scale vertically inserted on the right) but are located in the very tip of the ceramic backbone (see also the tip of the glass capillary). For the present experiments, the electrode/glass capillary assembly was inserted into the medial frontal cortex of mice. **f and g** show a coronal section of the frontal cortex (Prl, prelimbic cortex) and dorsomedial striatum, respectively, and the approximate placements and relative dimension of the assembly.

The construction of choline sensors and the amperometric measurement scheme were detailed previously (references above). Briefly, four platinum-iridium (Pt/Ir) recording sites (15 x 333 μm) were arranged in two pairs positioned linearly along the shank of the ceramic backbone probe (Quanteon; Nicholasville, KY). The members of a pair of recording sites were separated by 30 μm and pairs were separated along the vertical axis by 100 μm (Fig. 1).

An exclusion layer of *m*-(1,3)-phenylenediamine (mPD) was electroplated onto the surface of each recording site by applying a constant voltage of 0.85 V versus an Ag/AgCl reference electrode (Bioanalytical Systems; West Lafayette, IN) for 5 min to prevent sensing of electroactive interferents such as ascorbic acid (AA) and catecholamines. After allowing 24 hours for sites to dry, the top pair of electrodes, further named choline sensors, were coated with a solution of choline oxidase (Sigma-Aldrich; St. Louis, MO) cross-linked with a BSA and glutaraldehyde mixture by manual application of microdroplets using a 1-μl Hamilton syringe. The lower pair of sites were coated with only the BSA and glutaraldehyde mixture to identify currents unrelated to the presence of choline (hence named “sentinels”). Following a 48-hr minimum incubation period to allow for optimal adherence of the enzyme layer, recording sites were calibrated to determine the sensitivity for choline, selectivity for choline versus interferents, and limit of detection of choline. Calibrations were performed using a FAST-16 electrochemical system (Quanteon) to apply a constant voltage of 0.7 V versus an Ag/AgCl reference electrode in a heated bath of 40 mL 0.05 M PBS. After recording a 30-min baseline, 20 mM AA (interferent), 20 mM choline (analyte), and 2 mM dopamine were added to the bath to generate the following concentrations: 250 μM AA, 20, 40, and 60 μM choline, and 2 μM dopamine. Changes in amperometric current recorded at the individual electrode sites following the addition of each solution were used to calculate site sensitivity, selectivity, and limit of detection. In order to qualify for use in *in vivo* experiments, choline sensors were required to have a sensitivity of >5 pA/μM, choline:AA selectivity ratio >80:1, and a limit of detection <0.9 μM choline.

Probes that passed the above calibration criteria were prepared for *in vivo* experiments. Glass capillaries (1.0 × 0.58 mm, with filament; A-M Systems; Sequim WA) were pulled using a micropipette puller (Model 51210, Stoelting), and the tips were clipped until the inner diameter was approximately 15 μm. Pipettes were immobilized to the probe surface using clay and wax such that the tip of the pipette was positioned on the mid-line between the upper and lower site pairs, approximately 50 μm from the probe surface (Fig. 1e-g). Micropipettes were filled with one of the following solutions immediately prior to surgery: 5 mM choline, 5 mM choline and 10 μM HC-3, 20 μg/μL lidocaine HCl, or either 70 or 120 mM KCl. All drug solutions were prepared in DI water.

#### In vivo amperometric recordings

Mice were anesthetized using isoflurane gas (4% induction and 1-2% maintenance) and mounted in a stereotaxic frame on an isothermal heating pad to maintain a 37°C body temperature. Eyes were lubricated with ophthalmic ointment. A craniotomy and durotomy were performed above the right medial frontal cortex (AP: +1.9 mm; ML: −0.5 mm from bregma), or above the right dorsomedial striatum (AP: +0.50 mm; ML: −2.00 mm from bregma), using a bent 27-gauge needle for dura removal. The prepared microelectrode probe was then lowered 2.0 mm or 2.5 mm dorsoventrally from the top of the brain into cortex or striatum, respectively, while an Ag/AgCl reference electrode was implanted at a remote site in the contralateral hemisphere. Amperometric recordings were achieved using a FAST-16 electrochemical system and digitized at a sampling rate of 10 Hz. Pressure ejections began following a 60-minute baseline period. Choline sensing was accomplished through choline oxidase catalyzation of the conversion of choline into hydrogen peroxide and glycine betaine. The hydrogen peroxide byproduct was oxidized electrochemically via FAST application of 0.7 V versus the Ag/AgCl reference electrode. The resulting current, over background activity, was previously demonstrated to reflect either choline derived from newly released ACh and hydrolyzed by endogenous AChE (references above), or choline concentrations resulting from pressure ejections of choline into the recording area. In the latter case, diminishing choline currents were shown to largely reflect clearance by the CHT, as indicated by the attenuation of clearance efficiency following CHT blockade with HC-3 (Parikh et al., 2013; Parikh and Sarter, 2006; Parikh et al., 2006).

#### Choline clearance

We determined the impact of the Val89 variant on CHT-mediated clearance of choline pressure-ejected into the recording field. Micropipettes were filled with a 5 mM choline solution and pressure ejections of 10-35 nL (corresponding to a recorded amplitude of 10-35 μM choline) were administered at least one minute apart to limit inter-trial interference on choline clearance. The first 3 current traces within the 10-35 μM range were used for analysis.

In separate mice, exogenous choline clearance in the presence of HC-3 (Sigma-Aldrich) was determined to test the relative contribution of heterozygous and homozygous CHT Val89 expression, as compared to the expression of the CHT Ile89, to choline clearance. Micropipettes were filled with a cocktail of 5 mM choline and 10 μM HC-3. Pressure ejections were separated by a minimum of two minutes, and the first 3 ejection traces with a peak amplitude between 10-35 μM were used in subsequent analyses to avoid aggregating effects of serial HC-3 ejections. Clearance data from previous sessions in which only 5 mM choline was ejected were used as a baseline measure of comparison for all analyses regarding the inhibitory effects of HC-3.

In a third experiment, to quantify total voltage-dependent choline clearance, exogenous choline clearance was tested before and after administrating the voltage-gated sodium channel blocker, lidocaine HCl (Hospira Inc.; Lake Forest, IL). In addition to inhibiting neuronal activity (Tehovnik and Sommer, 1997; Malpeli and Schiller, 1979) that supports CHT surface trafficking (Ferguson et al., 2003), lidocaine-induced increases in sodium concentrations (Onizuka et al., 2008) and intracellular proton levels disrupt CHT function (Chu and Lee, 1994; Iwamoto et al., 2006). For this experiment, two micropipettes, one filled with 5 mM choline and the other filled with 20 μg/μL lidocaine HCl, were immobilized to the probe. Recording sessions consisted of 2 baseline pressure ejections of 5 mM choline, followed by a single 0.5 μL infusion of lidocaine HCl given over three minutes, a 2-minute rest period for drug diffusion, and finally, a second series of 2 post-lidocaine choline ejections (Malpeli and Schiller, 1979; see timeline in Fig. 5c). These parameters were also used in an experiment designed to verify the efficacy of lidocaine by measuring effects on depolarization-induced release of endogenous ACh. In one male WT mouse, pre- and post-lidocaine pressure ejections of KCl (70 mM; 40-60 nL) were used to generate ACh release events.

#### ACh release capacity following repeated depolarization

A final amperometric experiment determined the impact of the Val89 variant on ACh release following repeated depolarization. Repeated series of pressure ejections of potassium chloride (KCl; 120 mM; 30-40 nL per pressure ejection) were used to induce low-frequency, behaviorally-relevant episodes of ACh release. Choline currents following KCl-induced depolarization have previously been shown to reflect the hydrolysis of newly released ACh (Parikh and Sarter, 2006; Parikh et al., 2004; see Results for further justification and additional references). Each depolarization series consisted of 5 evenly distributed KCl ejections delivered over 2 min, with an ejection delivered every 24 s. Two depolarization series, separated by 15-20 min, were followed by a 5- and a 10-min pause, after which a single KCl-induced ACh release was measured (see timeline in Fig. 5a).

#### Analysis of choline currents

Choline currents were calculated by subtracting the background current recorded on sentinel sites from the choline-specific current recorded on choline sensors. Current was then converted to choline concentration based on calibration data. Choline clearance capacity was analyzed by calculating the slope of current decline from 40% to 80% of the peak choline concentration (Slope_40-80_). Slope_40-80_ was previously shown to be maximally sensitive to CHT inhibition by HC-3, indicating that this measure primarily reflects choline clearance via CHT-mediated transport (Parikh and Sarter, 2006). To enhance the illustration of the effects of HC-3 and lidocaine across the three genotypes, choline clearance traces were normalized by dividing individual values by the peak concentration obtained from the individual trace. Statistical analyses of clearance rates (Slope_40-80_) were always based on values extracted from raw traces.

### Determination of synaptosomal CHT density

#### Synaptosome preparation

Urethane-anesthetized mice were decapitated, and frontal cortices were dissected on ice immediately. Isolated tissues were homogenized in ice-cold 0.32 M sucrose and centrifuged at 1000 × *g* for 4 min at 4°C to remove cellular debris. Next, the supernatant was centrifuged at 12,500 x g for 15 min to yield a crude synaptosomal pellet. Pellets were stored at −80°C until fractioned.

#### Isolation and quantification of plasma membrane-enriched fractions

In addition to the determination of CHT density in total synaptosomal lysates, subcellular fractionation was conducted to isolate CHTs in a plasma membrane-enriched fraction (Parikh et al., 2013; Ferguson et al., 2003; Parikh et al., 2006). Briefly, the synaptosomal pellet was lysed in 5 mm HEPES-NaOH, pH 7.4, containing a cocktail of protease inhibitors (1.0 μg/ml leupeptin, 1.0 μg/ml aprotinin, 1.0 μg/ml pepstatin, and 250 μg/ml phenylmethylsulfonyl fluoride). The fraction was collected by spinning the lysate at 15,000 × *g* for 20 min. Proteins were extracted from each fraction with 5 mm HEPES-KOH (pH 7.2) solution containing 1% SDS, 1 mm EDTA, 1 mm EGTA, and protease inhibitor cocktail. Protein concentrations were determined by using a modified Lowry Protein Assay (Pierce). Equal quantities (25 μg) of protein from each fraction were subjected to immunoblot analysis. Proteins were separated on 4-15% Tris HCl polyacrylamide gels and transferred to PVDF membranes. Immunodetection of CHT bands was accomplished by incubating the membraned overnight with 1:2000 diluted rabbit anti-CHT polyclonal antibody (ABN458; Millipore). The membranes were then exposed to peroxidase-conjugated anti-rabbit secondary antibody and SuperSignal West Femto Maximum Sensitivity Substrate (ThermoFisher Scientific). The resulting chemiluminescent signal was acquired with a Chemidoc Touch Imaging System (Bio-Rad). Densitometric analysis of CHT-immunoreactive bands was performed by calculating the integrated pixel densities using NIH ImageJ software. The membranes were stripped for the detection of β-actin in all samples that served as a control to accommodate any differences in the sample loading during gel electrophoresis. CHT densities were normalized to the levels of β-actin-immunoreactive bands for each sample analyzed. Total synaptosomal CHT density was determined in frontal cortical tissue from 6 mice (3 females) per genotype. CHT density in the plasma membrane-enriched fraction was measured in tissues from 10 mice (5 females) per genotype.

### Disruptor-challenged attentional performance

#### Subjects

A total of N=45 mice, aged 12 weeks at the beginning of experiments, began acquisition training of the task. The final analyses were based on mice that reached asymptotic performance in the unchallenged version of the task and completed an additional sequence of sessions that included three sessions during which the disruptor was presented.

#### Water restrictions and body weights

Training and testing took place Monday-Friday between 11 am and 3 pm. Daily training of this task was previously shown to invoke a diurnal activity pattern in rodents (Gritton et al., 2013; Gritton et al., 2012; Paolone et al., 2012). Mice were weighed weekly. Prior to water deprivation, body weights ranged from 20-29 g (mean: 24.96 g). Beginning 5 days prior to the onset of training, water access was progressively reduced from *ad libitum* to 4 min per day. Water restrictions were adjusted individually throughout the experiment to maintain 90% of the mice’s original body weight, resulting in 3-6 min access to water after completion of the session. During weekends, mice were given unrestricted water access until Sunday 12 pm.

#### Task training and testing

The continuous signal detection task was previously characterized for the assessment of sustained attention in rats and humans (Demeter et al., 2008; McGaughy and Sarter, 1995) and subsequently adopted for testing mice (St Peters et al., 2011). Training and testing took place in eight operant chambers equipped with two center panel lights – the bottom light serving as the signal source -, a water dispenser, two retractable nose poke devices designed and built in-house (Michigan Controlled Access Response Ports, MICARPS; St Peters et al., 2011), and house light panel lights, a water dispenser, two moveable nose poke devices designed and built in-house (Michigan Controlled Access Response Ports, MICARPS; St Peters et al., 2011), and a house light (for technical specifications, including chamber dimensions, exact position and types of lights, illuminance measures, and the position of MICARPS, see St Peters et al., 2011).

Mice were trained to perform this task using the following steps. 1. Mice were initially trained to retrieve water from the water port (water was dispensed every 30 s for a single 20-minute session). 2, Both MICARPs were extended into the operant chamber and remained extended throughout the entire 40-minute session. All nose-pokes resulted in the reward of water (6 μL per reward). To pass this stage, mice were required to generate 20 rewards per session for three consecutive sessions. 3. Both MICARPs extended into the operant chamber and remained extended for 20 seconds, or until a nose-poke triggered retraction of both, with an intertrial interval (ITI) of 20 seconds. To pass this stage, mice must have retrieved 20 rewards per session for three consecutive sessions. 4. Both MICARPs extended into the operant chamber and remained extended for 4 s, or until a nose-poke triggered retraction, with an ITI of 12±3 s (criterion as in 3). 5. Mice were trained to discriminate signals (illumination of central panel light) from non-signals (no illumination) trials. Half of the mice were trained to report the presence of a signal with a left nose-poke, while the other half with a right nose-poke. Each 40-min session consisted of 160 trials (80 signals and 80 non-signals). One second after an event (signal or non-signal), MICARPs were extended into the chamber. In signal trials, the light remained illuminated until a nose-poke occurred or for 4 s (omission). Correct responses (hits, correct rejections) trigger water delivery, incorrect responses (misses, false alarms) triggered the onset of the ITI (see task illustration in Fig. 7a). This training stage also included forced correction trials. An incorrect response forced repetition of the trial. Three consecutive errors triggered a trial in which only the correct MICARP was extended, to suppress the manifestation of a side bias. To pass this stage, mice were required to score >60% hits and correct rejections. 6. Signal duration was shortened to 1 s, and the delay between signal/non-signal onset and MICARP extension was reduced to 0.5 s (criterion as in 5). 7. Correction trials were removed, and variable signal durations were implemented (500, 50, 25 ms) to prevent the manifestation of a fixed discrimination threshold (pass criterion: >60% hits to longest signals, >60% correct rejections, and <33% omissions for three consecutive sessions). 8. The house light in the back panel of the chamber was illuminated throughout the session. This is a crucial step for assessing sustained attention as it requires mice to continuously orient to and monitor the signal source (criterion as in 7).

#### Visual disruptor

Presentation of a visual disruptor disrupts performance and causes reward loss, thereby potentially triggering the implementation of top-down attentional control mechanisms to stabilize and recover performance (Demeter and Woldorff, 2016; Demeter et al., 2013; Demeter et al., 2008). Performance of the 40-min sessions was analyzed based on five 8-min trial blocks, and the disruptor (house light and panel light above the signal light flashing asynchronous at 0.5 Hz) was presented during the 2^nd^ and 3^rd^ 8-min block. Following a disruptor session, mice were required to regain unchallenged criterion performance for three consecutive sessions before being eligible for additional disruptor sessions. Sessions in which pre-disruptor performance reached chance level were excluded from final analysis. Neither the number of disruptor sessions excluded in the final analyses (median: 1.00, range: 3.00; Kruskal-Wallis ANOVA; H(2)=0.84, *P*=0.66) nor the number of sessions included (median: 2.00, range: 4.00; H(2)=0.34, *P*=0.95) differed by genotype.

#### Measures of performance

The percentage of hits, misses, correct rejections, false alarms, and omissions, for each 8-min trial block, were calculated. Hit and false alarm rates were used to compute signal detection theory-derived measures of perceptual sensitivity (*d*’) and response bias (*b*”_D_) (Green and Swets, 1974; Donaldson, 1992), as detailed previously (e.g., Demeter et al., 2008; Phillips and Sarter, 2020). The bias scores range from −1.0 to +1.0 with a score of zero depicting no bias, negative scores a liberal bias that is, a bias toward reporting the presence of a signal, and a positive score indicating a more conservative bias, that is, a bias toward reporting the absence of a signal.

### Structural modeling of the mSLC5A7 WT and I89V substitution

To identify a suitable structural template for mouse CHT we used Hidden Markov model profiles (HMM) as a descriptor for the transporter’s amino acid sequence. The HMM profile obtained after a three-iteration sequence scanning performed using UniRef30 sequence database was subsequently scanned against the HMMs of the sequences corresponding to each of the Protein Data Bank (PDB) X-ray structures (pdb70 database) as per standard procedure in HHpred server (Hildebrand et al., 2009; Zimmermann et al., 2018). This method has been shown to increase the successful identification of templates by 30%, especially for distantly related sequences. The *Vibrio parahaemolyticus* sodium/sugar symporter vSGLT (PDB id: 3DH4) (Faham et al., 2008) had the highest coverage (422 residues), the highest sequence identity (19%) and better correspondence between secondary structural elements and, as consequence, was selected as template during the structural modelling procedure. To further determine the architectural similarity of CHT and vSGLT we analyzed their hydrophobicity profiles. The hydrophobicity profiles for these proteins were constructed and aligned using AlignMe server (Stamm et al., 2014, 2013; Khafizov et al., 2010) and applying the Hessa et al. (2005) hydrophobicity scale and a 13-residue long window.

The initial sequence alignment between vSGLT and CHT obtained from HHpred was modified to align the first transmembrane helix (TM1) of vSGLT with that predicted in CHT by Topcons server (Tsirigos et al., 2015). The resulted alignment was then refined in an iterative process that used conservation scores obtained with Consurf server (Ashkenazy et al., 2016) as a guide. This procedure positions the most conserved residues packing towards the core of the protein and removes gaps within secondary structural elements when needed. The final alignment was used during the modelling production run where 2000 modelling iterations were performed with MODELLER (Webb and Sali, 2016). The model with the highest MODELLER probability distribution function score (molPDF), best ProQM score (Ray et al., 2010) and Procheck analysis (Laskowski et al., 1993) was selected as the final WT CHT model. A similar procedure was subsequently used to obtain the structural model of CHT Val89.

### Experimental design and statistical analysis

Choline clearance capacity was assessed using the rate (μM/s) of choline concentration decline from 40 percent to 80 percent of the choline transient peak (Slope_40-80_). An initial ANOVA of the effects of genotype was conducted to demonstrate that pressure ejections of choline produced similar peak amplitudes in the three strains. A two-factor mixed model ANOVA was used to determine genotype differences, and the effects of repeated (3) pressure ejections of choline, on choline clearance rate for choline-only ejection experiments. Two-way ANOVAs were used to test the effect of CHT inhibitor, HC-3, and lidocaine administration, on choline clearance rates (Slope_40-80_).

The effects of repeated depolarizations on evoked ACh release were analyzed using one-way and mixed model ANOVAs (see Fig. 5a for experimental manipulations and timeline). In each case, results from Shapiro-Wilks tests of Normality and Levene’s equality of variance supported the use of parametric analyses. Effects of the very first pressure ejection of potassium on absolute peak amplitudes of choline currents in the three strains were analyzed using one-way ANOVA. The effects of subsequent KCl ejections were expressed as a ratio of the effects of the first ejection. The effects of genotype on the peak amplitudes of currents evoked by subsequent depolarizations (2^nd^ to 5^th^ depolarization) were analyzed first by mixed model ANOVAs on the effects of genotype and KCl series (first versus second) separately for the 2^nd^ and 5^th^ KCl pressure ejection (with alpha set at 0.05/2). Based on the absence of significant effects from these two analyses, individual means over the two KCl series were analyzed for the effects of genotype and KCl ejection number (2^nd^ versus 5^th^). Peak amplitudes of currents recorded during the recovery tests, 5 and 10 min after the depolarization series, were expressed relative to the peak amplitudes of the first KCl-induced choline current of that series. Effects of genotype and series were analyzed using mixed model ANOVA, once again supported by the results of Shapiro-Wilk and Levene’s equality tests. CHT densities in total synaptosomal lysates and plasma membrane-enriched fractions were compared between the genotypes using one-way ANOVA. To determine effects on the performance in the continuous signal detection task, signal detection theory-derived parameters of perceptual sensitivity and bias were computed (see above). Mixed model ANOVAs determined effects of genotype and signal duration on unchallenged performance, and effects of trial block and genotype on the performance in the presence of a disruptor. To determine changes in bias in response to the disruptor, the difference between scores obtained during disruptor and post-disruptor trial blocks (2-5) were subtracted from scores indicating bias during the first pre-disruptor trial block. For each mouse, these differences were averaged across disruptor sessions and analyzed to determine effects of genotype and trial block using ANOVA. Finally, effects of task block and genotype on errors of omissions were determined for both task conditions.

For parametric mixed-model ANOVAs, in case of violation of the sphericity assumption, Huynh-Feldt-corrected F values, along with uncorrected degrees of freedom, are given. Tests used for *post hoc* multiple comparisons are identified in figure legends. Data graphs depict individual values, means and 95% Confidence Intervals (CI). Statistical analyses were performed using SPSS for Windows (version 17.0; SPSS) and GraphPad Prism Version 8.2.1. with alpha set to 0.05. Exact *P* values were reported (Greenwald et al., 1996; Sarter and Fritschy, 2008; Michel et al., 2020). For major results derived from parametric tests, effect sizes (Cohen’s *d*) were indicated (Cohen, 1988).

## Results

### Impact of Val89 on CHT-mediated choline clearance

#### Reduced clearance of exogenous choline

These experiments were designed to determine, *in vivo,* the capacity of the CHT in the cortex and striatum of CHT I/I, I/V and V/V genotypes (see Figure 1 for an illustration of the measurement scheme and the placement of the electrode/glass capillary in cortex and striatum). The clearance of exogenous choline was determined by analyzing the component of the clearance curve, Slope_40-80_, that was previously demonstrated to reflect primarily CHT-mediated transport (see Methods for details and references).

Following pressure ejections of 5 mM choline into the prelimbic recording region, peak choline currents were reached within 1-2 s, followed by rapid decreases in current (Fig. 2a). Peak currents did not differ between genotypes (F(2,36)=1.58, *P*=0.22; Fig. 2b; note that while currents were recorded at 10 Hz, and analyses were based on 10-Hz data, for clarity the figures show data at a 2 Hz sampling rate).

**Figure 2.**
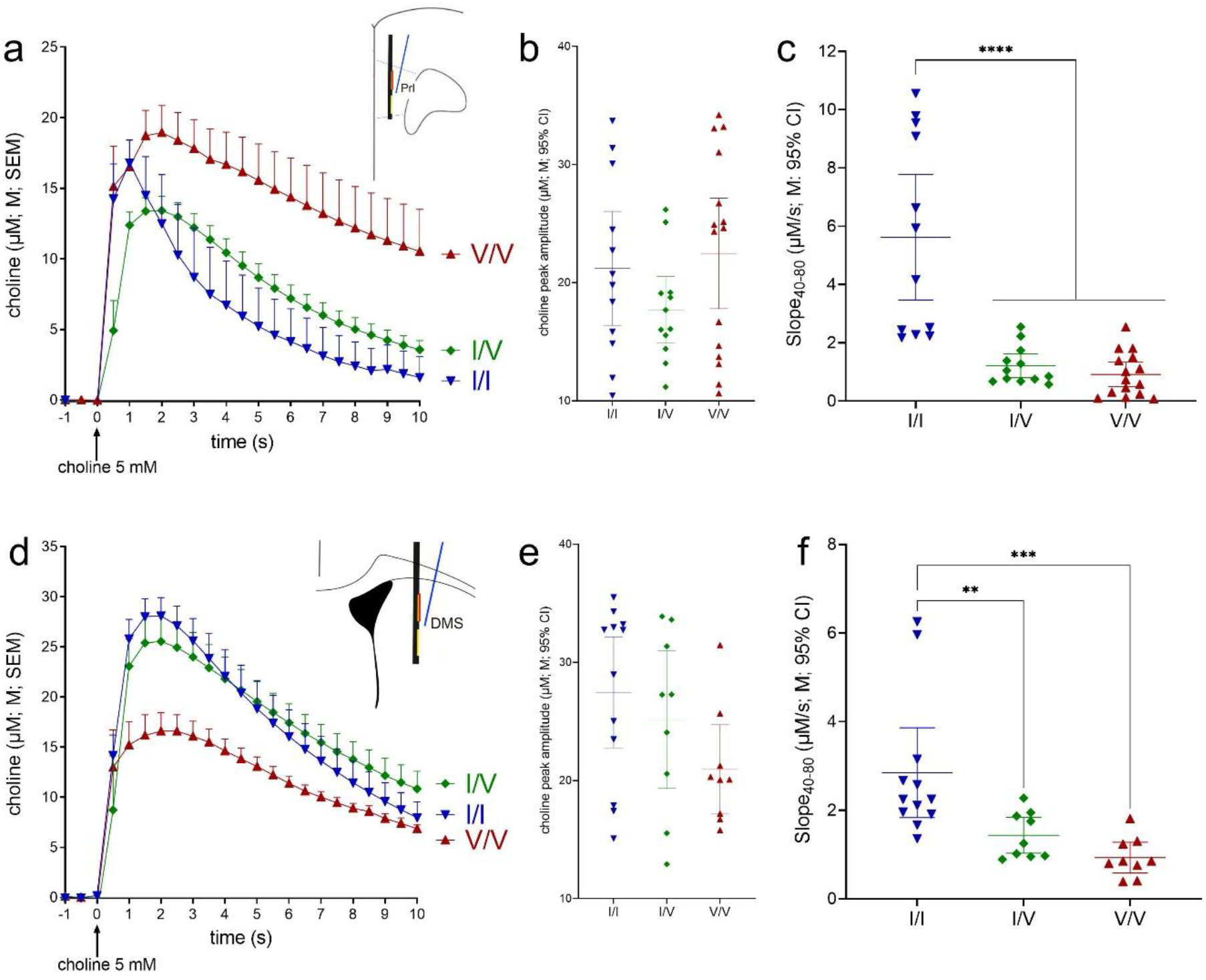
Impact of CHT Val89 on clearance of exogenous choline in mouse prelimbic cortex (a-c) and dorsomedial striatum (d-f; for illustrative clarity, choline currents, recorded at 10 Hz, are shown at 2 Hz). Following insertion of the electrode and glass capillary into the brain (see Fig. 1) and a 60-min baseline period, repeated pressure ejections of 5 mM choline (10-35 nL) were given at 1-2 min intervals. **a** Choline ejections occurred at time 0. **b** Peak choline currents did not differ across genotypes (ANOVAs are described in Results). **c** Cortical choline clearance rates were calculated for the periods of current decay from 40% to 80% (Slope_40-80_) of peak amplitudes (μM/s; shown are individual values, means and 95% CI; WT, inverted blue triangles, 12 ejections from 4 mice; CHT I/V, green diamonds, 12 ejections from 4 mice; CHT V/V, red triangles, 15 ejections from 5 mice). Clearance rates in both I89V-expressing strains were markedly reduced relative to WT mice (Least Significant Difference (LSD) test). Choline clearance in the striatum (**d-f**) likewise was significantly suppressed in I89V-expressing mice (WT: 4 mice with 3 ejections each; CHT I/V and CHT V/V mice: 3 mice with 3 ejections each). As averaged peak levels in **d** suggested relatively lower peak currents in Val89 mice, contrasting with the absence of differences between absolute peak currents (**e**), an additional analysis revealed a significantly longer time period over which peak currents were distributed in Val89 mice when compared with WT mice (Results), leading to relatively lower averages shown in **d** (see Results; note that, in **f**, the main genotype effect and multiple comparisons remain significant after removal of the two highest I/I data points; stars in **c** and **f** and in subsequent figures depict results from *post hoc* multiple comparisons; .*,**,***,****: *P*<0.05. 0.01, 0.001, 0.0001, respectively).

A mixed-model ANOVA on the effects of genotype and repeated pressure ejections on normalized choline clearance (Slope_40-80_) indicated a main effect of genotype (F(2,36)=6.91, *P*=0.01, Cohen’s *d*=2.35) and neither an effect of repeated pressure ejections of choline nor a significant interaction between the two factors (main effect: F(2,36)=0.22; interaction: F(4,36)=0.62; both *P*>0.41). Multiple comparisons (LSD) indicated significantly slower decay rates in CHT I/V and CHT V/V mice when compared with WT mice (*P*<0.012), but no gene dosage effect (*P*=0.83). Figure 2c illustrates that decay rates in CHT Val89 mice on average were reduced by over 77% (CHT I/V) and 83% (CHT V/V) relative to WT mice. The low residual decay rates in CHT I/V (25^th^ percentile: 0.7 μM/s; 75^th^: 1.6 μM/s) may have prevented the demonstration of even lower decay rates in CHT V/V mice. This issue was further addressed by the experiments which assessed effects of HC-3 and lidocaine (below).

Similarly, striatal choline clearance rates in CHT Val89 mice were significantly lower than in WT mice (F(2,27)=8.94, *P*=0.001, d=1.63). Furthermore, as also was the case in cortex, multiple comparisons (Fig. 2f) did not indicate a significant gene dose effect.

Following pressure ejections of choline into the dorsomedial striatum, inspection of the averaged choline current traces (Fig. 2d) suggested lower peak currents in CHT V/V than in WT and I/V mice. However, peak amplitudes did not differ across genotypes (F(2,27)=2.37, *P*=0.11; Fig. 2e). The averaged traces shown in Fig. 2d suggested longer lasting peak levels in CHT Val 89 mice when compared with WT mice. We therefore conducted a *post hoc* analysis of the timing of peak currents. Currents peaked between 1.8 and 2.1 s after pressure ejections and peak times did not differ across genotypes (F(2,27)=0.39, *P*=0.68). However, peak times were considerably more variable in CHT Val 89 when compared with WT mice (SD: WT: 0.38 s I/V: 0.98 s; WV/V: 1.46 s). Multiple F-tests for the Equality of Two Variances (at α/3) confirmed that peak current variances in CHT Val89 mice were significantly larger than in WT mice (WT vs. I/V: F(11,8)=6.70, *P*=0.005; WT vs V/V: F(11,8)=14.97, *P*<0.001; I/V versus V/V: F(8,8)=2.23, *P*=0.28). Thus, the seemingly lower levels of averaged peak currents in CHT Val89 mice seen in Fig. 2d did not reflect lower peak levels (Fig. 2e), but that the timing of current peaks was distributed over significantly longer periods of time, therefore lowering averaged values. The reason for this effect of genotype in the striatum is unknown. Because of our focus on CHT capacity in the cortex (see Introduction), subsequent experiments assessed effects solely in that brain region.

#### Impact of hemicholinium-3

We co-pressure ejected choline and the competitive CHT inhibitor HC-3 to demonstrate that the choline clearance measure used above (Slope_40-80_) indeed reflects primarily CHT-mediated choline clearance (Parikh and Sarter, 2006), and to explore whether CHT-mediated choline transport can be further reduced in Val89 mice by pharmacological blocking of the transporter (see Materials and Methods for details about the ejection parameters). Fig. 3a illustrates the marked slowing of choline clearance in the presence of HC-3 in WT CHT I/I mice, contrasting with the absence of a change in the slopes of the decay curves in CHT I/V (Fig. 3b) and CHTV/V (Fig. 3c) mice. The resulting Slope_40-80_ values are shown in Fig. 3d.

**Figure 3.**
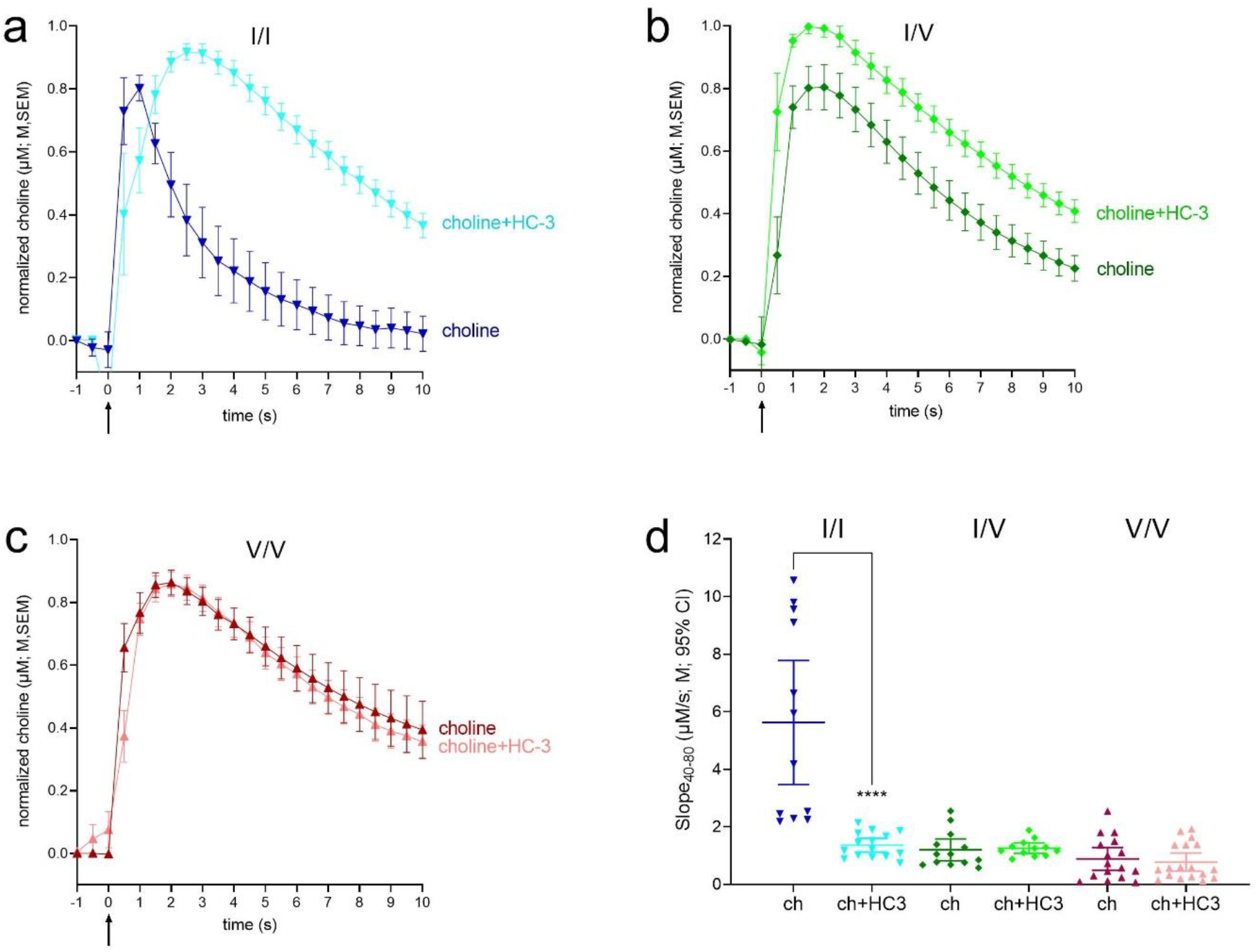
Impact of the CHT competitive inhibitor HC-3 on choline clearance in WT and CHT Val89 mice. **a-c** Averaged traces of exogenous choline clearance with and without HC-3 (for visual clarity, traces shown here and in Fig. 4 were normalized to the peak current values). Traces and slopes in the absence of HC-3 were obtained from a separate group of mice (Fig. 2). Choline clearance in the presence of HC-3 was determined from 15 traces in WT (n=5), 12 traces in CHT I/V (n=4), and 18 traces in CHT V/V (n=6), with 3 traces selected from each mouse. Darker colored lines represent choline clearance traces following pressure ejections of 5 mM choline alone, while lighter colors show clearance following ejections of a solution containing 5 mM choline and10 μM HC-3, with each solution ejected at time 0. **d** Individual choline clearance rates (Slope_40-80_) following ejections of 5 mM choline (ch) only and a 5 mM choline and 10 μM HC-3 cocktail (ch+HC3). Only clearance rates recorded in WT mice were significantly slowed by HC-3. Following HC-3 ejections, residual clearance rates did not differ across genotypes (LSD test).

ANOVA of the effects of genotype and HC-3 on choline clearance (Slope_40-80_) indicated main effects and a significant interaction between the two factors (genotype: F(2,167)=83.46, *P*<0.0001, *d*=1.9; HC-3: F(1,167)=74.99, *P*<0.0001, *d*=1.3; interaction: F(2,167)=57.11, *P*< 0.0001, *d*=1.5). *Post hoc* multiple comparisons indicated a HC-3-induced significant decline in the clearance rate in WT mice (*P*<0.0001), but not in CHT I/V or CHT V/V mice (both *P*>0.81; Fig. 3d). Indeed, choline clearance rates did not differ significantly between the genotypes in the presence of HC-3 (all *P*>0.06). A final *post hoc* comparison between absolute peak choline currents recorded following pressure ejections of solutions including and not including HC-3 (t(25)=0.09, *P*=0.93; data not shown) confirmed that effects on choline clearance were not confounded by differences in the volume and constitution of the ejection solution.

Together, the findings from the experiments on the effects of HC-3 confirm that the measure of choline clearance (Slope_40-80_) reflects CHT-mediated clearance, and they indicate that the low clearance rates in both variants were not measurably reduced by HC-3-induced blockade. As the affinity of HC-3, in the absence of extracellular choline, is unaffected by the Val89 variant (Okuda et al., 2002), the unexpectedly large impact of both variants on choline clearance, indicated by residual clearance rates below 1 μM/s, may have limited the demonstration of any further inhibition by HC-3. However, as the CHT supports vital functions (Ferguson et al., 2004) and thus was expected to remain functional in Val89 mice, we conducted an experiment designed to demonstrate such residual functionality.

#### Inactivation by lidocaine

Inactivation of neuronal activity and suspension of CHT activity by lidocaine (Tehovnik and Sommer, 1997; Malpeli and Schiller, 1979; Iwamoto et al., 2006; Onizuka et al., 2008) was implemented to determine whether voltage-dependent choline clearance can be detected in CHT Val89 mice. In the presence of lidocaine, residual choline clearance rates are thought to reflect diffusion (Cass et al., 1993) and, potentially, choline uptake via low-affinity, electrogenic transporters other than CHT (Michel et al., 2006). Therefore, lidocaine was expected to suppress the already low choline clearance rates in Val89 mice.

We confirmed the efficacy of lidocaine to suspend neuronal depolarization in the recording area by measuring K^+^-evoked, depolarization-induced ACh release in one WT mouse. Three traces each prior and ~2-4 min after the lidocaine infusions (0.5 μL over 3 min) are illustrated in Fig. 4a. Peak amplitudes reached about 10 μM prior to inactivation and 2 μM following lidocaine (Fig. 4b), consistent with prior results showing less than complete inactivation following intracranial infusions of relatively small volumes of lidocaine (Tehovnik and Sommer, 1997).

**Figure 4.**
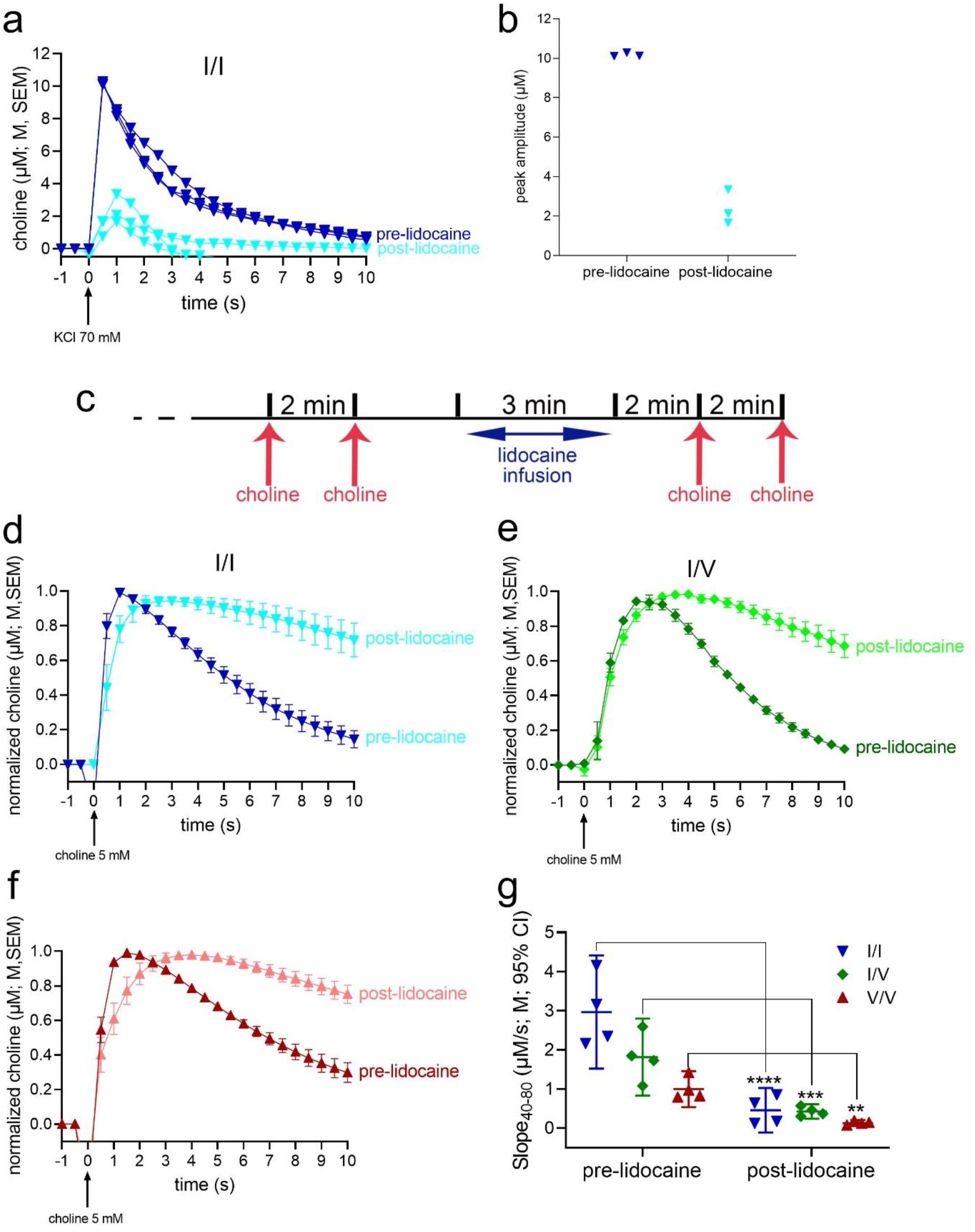
Reduced choline clearance rate following inactivation of neuronal activity and CHT function with lidocaine. An initial experiment, using one WT mouse, was conducted to verify the efficacy of lidocaine (0.5 μL of 20 μg/μL infused over a consistent rate of 0.17 μL /min) to inactivate the recording area. Prior to lidocaine infusions, KCl (70 mM) pressure ejections reliably generated ACh release events, peaking at about 10 μM (see dark blue traces in **a**). Two to three min following lidocaine infusions, peak choline currents were markedly reduced (**b**), yet not completely abolished, as would be expected from the effects of a relatively small volume of lidocaine (Tehovnik and Sommer, 1997). **c** Timeline of choline pressure ejections and lidocaine infusion used to test choline clearance in the three strains. Following a 60-min baseline period, two initial pressure ejections of choline (5 mM), separated by 2 min, were followed by the infusion of lidocaine and, 2 min later, by another two pressure ejections of choline. **d,e** and **f** represent averaged (over 8 traces from two mice per genotype) pre- and post-lidocaine currents evoked by exogenous choline in WT (**d**), CHT I/V (**e**) and CHT V/V (**f**) mice. **g** ANOVA indicated main effects of genotype and lidocaine as well as a significant interaction (Results). LSD multiple comparisons tests indicated significant reductions of choline clearance following inactivation in all 3 genotypes (**g**). The significant interaction between the effects of genotype and lidocaine also reflected that choline clearance rates no longer differed between the genotypes (all *P*>0.10).

Choline clearance rates were assessed following the timeline of choline pressure ejections and lidocaine infusions illustrated in Fig. 4c. First, we compared choline clearance rates (Slope_40-80_) measured prior to lidocaine infusions with those measured in our initial experiment (Fig. 2b). This analysis reproduced the main effect of genotype (F(2,45)=12.23, *P*<0.001), reflecting that CHT I/I clearance rates were higher than in CHT I/V (*P*=0.0004) or CHT V/V mice (*P*<0.0001). Furthermore, choline clearance rates across the three strains did not differ between the two experiments (F(1,45)=1.26, P=0.27).

As illustrated in Figs. 4d-g, lidocaine-induced inactivation reduced choline clearance in all three genotypes (main effect of lidocaine: F(1,27)=104.40, *P*<0.0001, *d*=3.9), with relatively greater effects seen in WT than in Val89 mice (main effect of genotype: F(2,27)=21.12, *P*<0.0001, *d*=2.5; interaction: F(2,27)=13.40, *P*<0.0001, *d*=2.0). Moreover, these results are consistent with an absence of genotype-related differences in clearance rates when transport by CHT and other electrogenic transporters is suppressed (all *P*>0.10; Fig. 4g).

### Attenuated ACh release capacity following repeated depolarization in vivo

The results described above indicate a highly significant reduction in choline clearance in CHT Val89 mice when compared with mice expressing the WT CHT Ile89 transporter. Such a reduction is expected to impact the capacity of cholinergic neurons to synthesize and release ACh, particularly when cholinergic activity is relatively high and presynaptic “reloading” of newly synthesized ACh is taxed. Basal forebrain cholinergic neurons exhibit highly diverse firing patterns, with firing bursts of subpopulations of these neurons reaching >80 Hz (e.g., Laszlovszky et al., 2020; Manns et al., 2000; Duque et al., 2000). However, given our focus on cholinergic activity during attention, we employed a markedly lower frequency and degree of cholinergic taxation to foster an interpretation of results in the context of the inattentive phenotype in humans expressing the Val89 variant. Attention tasks such as those used in humans and rodents to reveal attentional impairments typically consist of trial frequencies of ~ 1 to 0.1 Hz (e.g., Berry et al., 2014; Demeter et al., 2008). In such tasks, ACh release events occur during successful cue detection (Gritton et al., 2016). We tested ACh release capacity in CHT Val89 mice following a relatively small number of potassium-induced depolarizations, delivered at a relatively low rate (see Methods; Fig. 5a). ACh release capacity was assessed 5 and 10 min after such depolarization trains (Fig. 5a), mirroring recovery periods that previously revealed ACh release deficiencies in heterozygous CHT knockout mice (Parikh et al., 2013).

**Figure 5.**
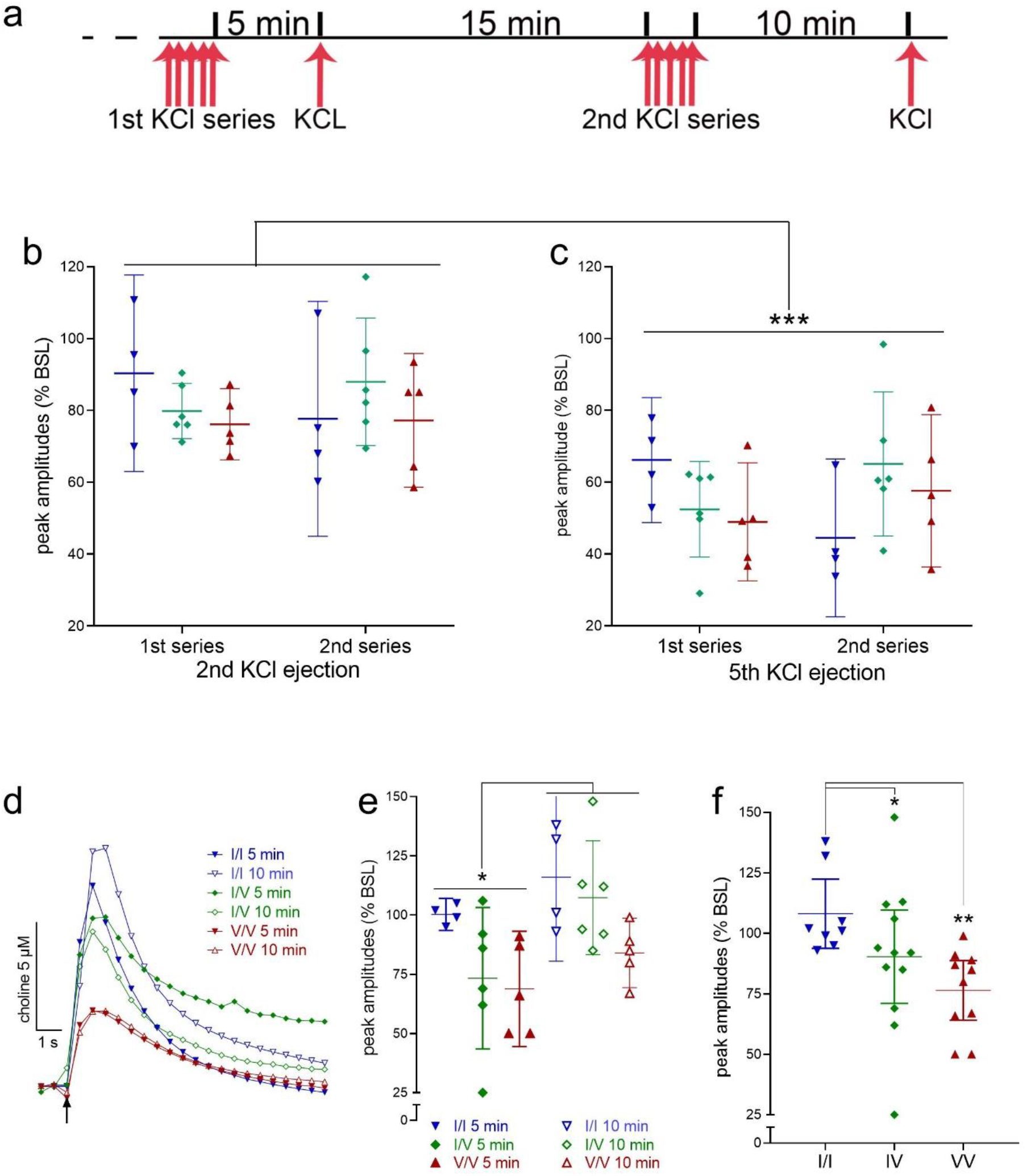
Impact of CHT Val89 on ACh release capacity following repeated depolarization. **a** Schematic illustration of the timeline of KCl pressure ejections. Following a period of calibration, an initial train of 5 KCl ejections were administered over 3 min and followed, 5 min later, by a first release capacity test. After a 15-min pause, a second series of depolarizations was followed, 10 min later, by a second release capacity test. Peak amplitude currents following the first KCl ejection of either series did not vary by genotype (not shown). **b,c** show the peak amplitudes of currents evoked by the 2nd and 5th ejection, respectively, of either series, expressed as a ratio of the amplitudes of the first ejection (n=4 WT, 6 CHT I/V, and 5 CHT V/V mice; lines and bars in **b,c,e,f** depict means and 95% CI). Compared with the 2nd depolarization, the 5th resulted in significantly smaller amplitudes across genotypes and the two series. **d** Representative traces of currents evoked at the 5-min and 10-min recovery tests, illustrating attenuated ACh release in CHT I/V and CHT V/V mice. **e** Peak amplitudes of the recovery currents, expressed as the percent of the amplitude of the first depolarization of the respective series, indicated significantly greater recovery after the longer 10-min pause compared with peak currents recorded 5-min after the depolarization train (see Results for ANOVA). The analysis of the effects of series and genotype also revealed a main effect of genotype, illustrated in **f**, that reflected significantly lower peak amplitudes in CHT I/V (12 ejections from n=6) and CHT V/V (10 ejections from n=5) mice relative to peak amplitudes in WT mice (8 ejections from n=4; LSD test; see **f** for *post hoc* multiple comparisons).

The very first pressure ejection of KCl produced statistically similar choline current peaks in the three genotypes (F(2,12)=0.79, *P*=0.48; M, SEM (μM choline): WT: 11.65±2.06; CHT I/V: 10.47±1.55; CHT V/V: 8.15±2.24; not shown). The peak amplitudes of the choline currents evoked by the subsequent 2^nd^, 3^rd^, 4^th^ and 5^th^ pressure ejections of KCl were expressed as percent of the peaks of the first ejection from the respective depolarization series. Irrespective of genotype and series, the repeated depolarizations during either series appeared to yield increasingly smaller peak amplitudes. Figs. 5b and c show the relative peak amplitudes evoked by the 2^nd^ and 5^th^ ejection of both series. There were no effects of genotype or KCl series (1^st^ versus 2^nd^; see Fig. 5a) on peak amplitudes evoked by the 2^nd^ or 5^th^ KCl ejections (main effects of genotype, series and interactions, analyzed separately for the 2^nd^ and 5^th^ ejection, with alpha=0.05/2: all F<2.53, all *P*>0.12). Therefore, the mean relative peak amplitude for each mouse was used to compare the effects of the 2^nd^ and 5^th^ ejection. This analysis indicated that irrespective of genotype, peak amplitudes were markedly reduced following the 5^th^ ejection (main effect of ejection: F(1,12)=88.89, *P*<0.001; Cohen’s *d*=5.08; overall M, SEM: 2^nd^ ejection: 81.54±2.56%; 5^th^: 56.03±2.11%; Figs 5b,c). These findings indicate the efficiency of the relatively low-frequency depolarization trains with KCl to reveal the limits of cholinergic terminals to synthesize and release ACh.

Five minutes following the first and 10 minutes following the second KCl-induced depolarization series, one additional depolarization was administered to test the “reloading” capacity of cholinergic neurons in the 3 genotypes. Representative traces of absolute KCl-induced currents (peaks matching the means of the genotype) are shown in Fig. 5d and suggested reduced amplitudes in CHT I/V and CHT V/V mice, relative to CHT I/I mice, at both time points. To indicate the degree of synthesis and release recovery, peak amplitudes of choline currents were expressed as a ratio to the peak amplitudes of currents evoked by the first depolarization from the respective series (Fig. 5e and f). Irrespective of genotype, relative peak amplitudes recorded after the 10-min pause were higher than those recorded after the 5-min pause (main effect: F(1,12)=6.11, *P*=0.03; Cohen’s *d*: 1.42; M, SEM: 5 min: 80.86±5.18%; 10 min: 102.38±5.69%; Fig. 5e). Moreover, a main effect of genotype (F(2,12)=7.16, *P*=0.009; Cohen’s *d*=2.18; Fig. 5f) reflected a relatively greater recovery of ACh release in CHT I/I mice than either of the CHT Val89-expressing mice (no difference between CHT I/V and CHT V/V; genotype x series: F(2,12)=0.56, *P*=0.59, *d*=0.61; multiple comparisons, using LSD, are shown in Fig. 5f). Taken together, this experiment indicated the efficiency of the depolarization series to reduce ACh output and an attenuated capacity of cholinergic neurons of CHT I/V and CHT V/V mice to regain baseline ACh release levels after relatively long, 5- and 10-min pauses.

### CHT densities do not differ between genotypes

Because of our overall focus on the role of fronto-cortical cholinergic signaling in attention (e.g., Howe et al., 2017; Gritton et al., 2016) and that CHT I/V humans fail to activate right frontal cortex during attention (Berry et al., 2015), frontal cortices were harvested for the analysis of CHT densities in total synaptosomal lysates and synaptosomal plasma membrane-enriched fractions. We previously demonstrated that CHT density in the latter relatively precisely predicts choline clearance capacity (Koshy Cherian et al., 2019; Parikh et al., 2013; Apparsundaram et al., 2005; Ferguson et al., 2003). As illustrated in Figure 6, CHT densities in neither total synaptosomal lysates nor plasma membrane-enriched fractions differed by genotype (both F<0.58, both *P*>0.56). These findings are consistent with the view that attenuated choline clearance and ACh release following repeated depolarizations likely reflects a conformational modification of the protein induced by the Val89 variant that impacts transporter function rather than translation, stability or surface trafficking of CHT protein.

**Figure 6.**
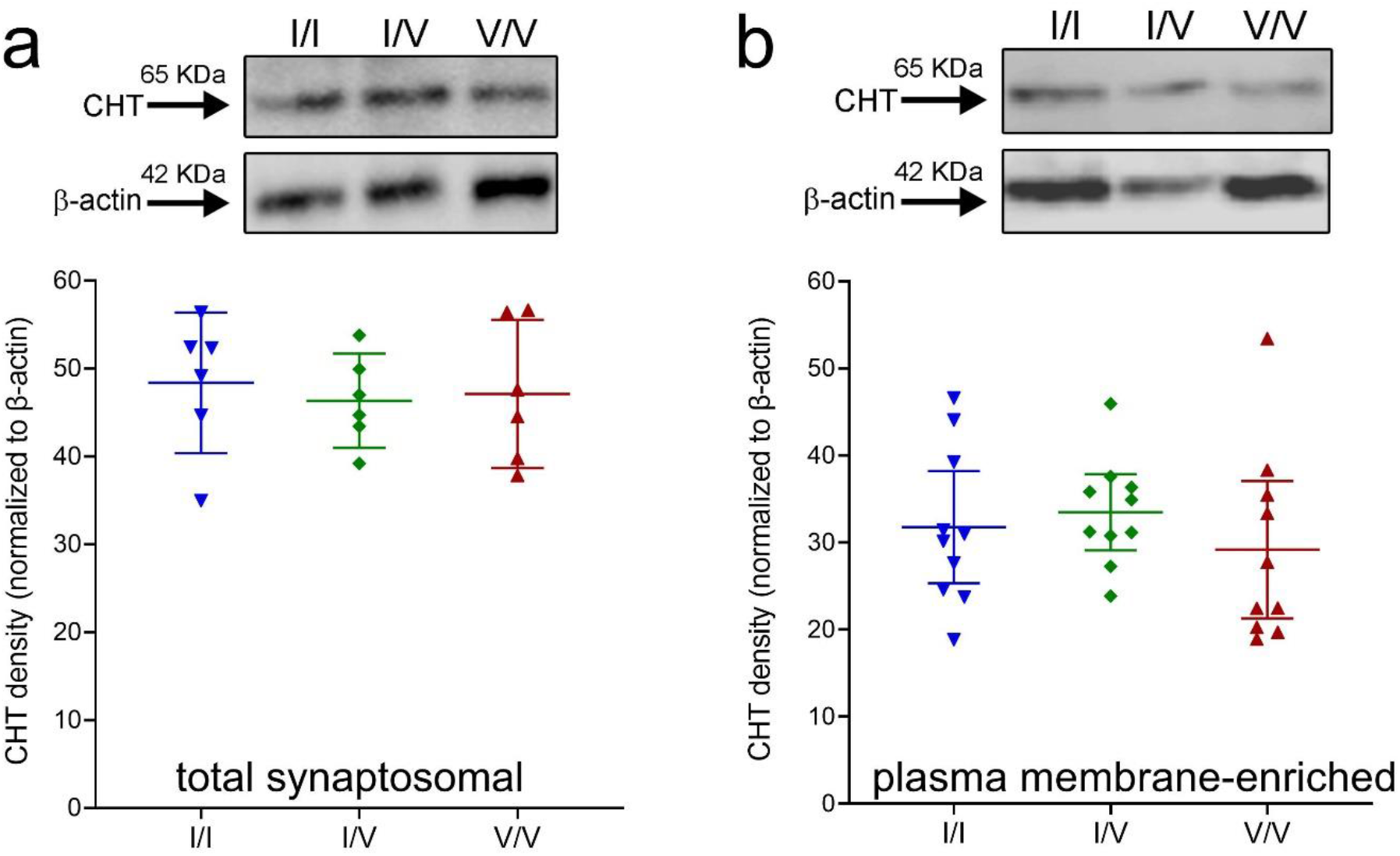
Top: Western blots showing the presence of the CHT in frontal cortical total synaptosomal lysates (**a**, top) and synaptosomal plasma-membrane-enriched fractions (**b**, top; normalized against β-actin; each lane is based on tissue from one individual; 25 μg protein loaded in each well; total and plasma membrane-enriched fractions were derived from separate groups of mice). CHT densities in neither total lysates (**a**) nor plasma membrane-enriched fractions (**b**) differed between the genotypes (graphs show individual values, means and 95% CI).

### Attentional control deficit in Val89 mice

Mice were trained to perform a continuous signal detection task that was previously utilized to measure sustained attention in rats and humans (Demeter et al., 2008) and modified for use in mice (St Peters et al., 2011). Subjects were trained to report the presence or absence of a signal in trials that began either with a signal (variable duration) or a blank. Responses were hits and misses, and correct rejections and false alarms, respectively, with hits and correct rejections yielding water reward (Fig. 7a). Hits and false alarms were used to compute signal detection theory-derived measures of perceptual sensitivity *(d’)* and response bias *(B’_D_*) (Green and Swets, 1974; Swets et al., 1961; Donaldson, 1992). Val89 mice were not expected to exhibit deficits in perceptual sensitivity (Berry et al., 2015), but to alter their bias in response to a performance challenge. This prediction was derived in part from evidence in unscreened humans who performed a human version of the present signal detection task and found to adopt a relatively more conservative criterion in response to the presentation of a visual disruptor. In other words, the disruptor evoked a relatively greater propensity to report the absence of a signal (Demeter et al., 2008), thereby maintaining and recovering above-chance levels of response accuracy and reward rates. Thus, WT mice were expected to respond to a visual disrupter (see Methods) by shifting to a more conservative bias *(B’_D_* scores between 0 and 1), whereas Val89 mice were predicted to not change their bias, reflecting that bottom-up, or signal-driven mechanisms predominantly supported their performance.

**Figure 7.**
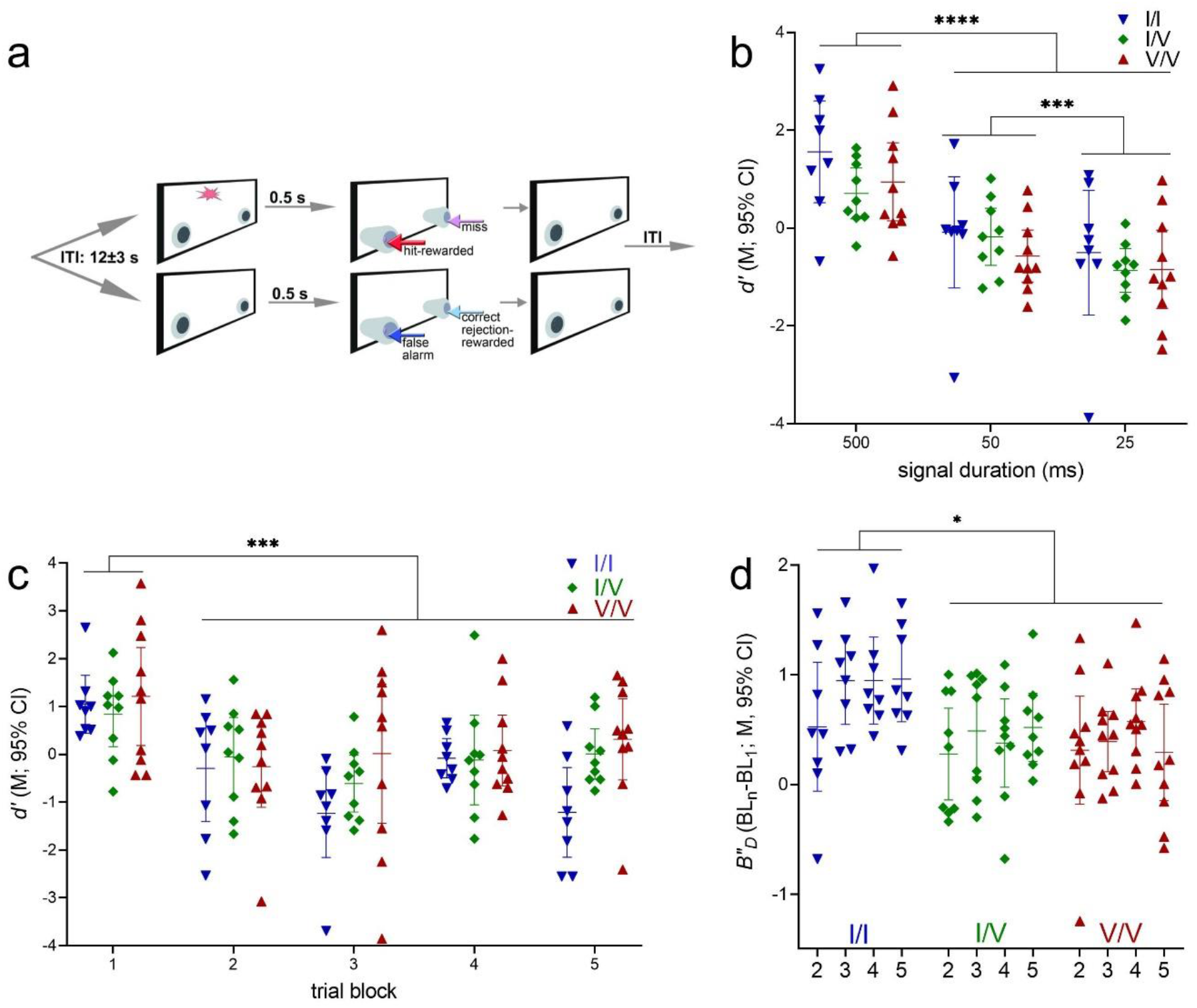
Performance of a continuous signal detection task and effects of a visual disruptor (WT: n=8, 3 females; CHT I/V: n=9, 4 females; CHT V/V: n=10, 2 females). **a** Illustration of the two trial types and main events of the signal detection task employed to assess sustained attention in mice. The task was previously cross-validated in rats and humans (Demeter et al., 2008) and adopted for testing mice (St Peters et al., 2011). Briefly, trials began either with a light signal of variable duration (top) or a blank (bottom), followed by the extension of nose-poking device into the chamber. Half of the mice were trained to nose-poke on the left to report the presence of a signal and on the right to report the absence of a signal. Correct responses (hits in signal trials and correct rejections in non-signal trials) were rewarded with water delivery while incorrect responses (misses and false alarms, respectively) triggered the onset if the intertrial interval (ITI; see Methods for more details). Signal detection theory-derived measures of perceptual sensitivity *d’* and response bias *b”d* were used to characterize effects of task parameters (signal duration, block of trials) and a visual disruptor. **b** As expected, asymptotic performance of the unchallenged version of the task revealed a main effect of signal duration on perceptual sensitivity but, similar to humans heterozygous for the Val89 variant (Berry et al., 2015), no genotype effects (**b-d** depict measures calculated over longest signal durations, as scores to shorter signals indicated chance level performance). **c** The visual disruptor (house lights flashing at 0.5 Hz during task blocks 2 and 3) was highly effective as indicated by near or at chance level performance of all mice during the remainder of the session. **d** During and after the disruptor challenge, WT mice, but not Val89 mice, adopted a more conservative criterion, that is, they were less likely to report the presence of a signal. Such a criterion shift to attentional challenges is thought to reflect the deployment of top-down attentional control mechanisms to support performance. This finding suggests that Val89 mice predominantly employed bottom-up, signal-driven mechanisms to perform this continuous signal detection task.

#### Unchallenged task performance

Upon reaching asymptotic performance in the unchallenged task, mice reliably reported the presence of signals in a signal duration-dependent manner (mean percent hits to 500, 50 and 25 ms signals: 72.39%; 47.05%; 38.45%) and accurately reported the absence of signals (mean percent correct rejections: 70.20%). As was expected given signal duration-dependent hit rates, perceptual sensitivity *d’* varied significantly by signal duration ((F(2,48)=146.40, *P*<0.001). However, there were no effects of genotype or interactions between genotype and signal duration on this measure (both F<2.80, all *P*>0.07; Fig. 7b).

As performance measures calculated over hit rates to 50 ms and 25 ms signals approached chancel level (Fig. 7b), *b”_D_* scores were computed only over hits to longest signal durations. Performing the unchallenged task, mice exhibited a mildly liberal bias (*b”_D_*=-0.15 (mean). *b”_D_* scores did not differ by genotype, and there were not interactions involving the factor genotype on bias (all F<1.44; all *P*>0.21; not shown).

The relative number of omissions remained relatively low (grand mean: 17.85%) but increased across blocks of trials from 9.31% in block 1 to 32.06% in block 5 (F(1.94, 46.58)=15.84, *P*<0.001). There were no significant effects of genotype and no interactions between genotype and block on omissions (both F<1.79, both *P*>0.18; not shown).

#### Bias changes evoked by a disruptor challenge

Performance was challenged using a visual distractor, presented in the 2^nd^ and 3^rd^ blocks of trials. The robust and genotype-unrelated efficacy of the disruptor was indicated by a significant reduction of perceptual sensitivity, calculated over hits to longest signals (main effect of block on *d’;* F(4,96)=8.47, *P*<0.0001; Fig. 7c). Perceptual sensitivity was not affected by genotype (main effects and interactions with block: both F<2.42, both *P*>0.11).

Similar to the response biases derived from unchallenged task performance, during the first (pre-disruptor) trial block all mice exhibited a relatively liberal response bias, that is, a propensity to report the presence of a signal (no effect of genotype: F(2,24)=1.71; *P*=0.21; *b”_D_* (M, SEM): −0.29, 0.09). As a result of the disrupter presentation WT, but not Val89, mice exhibited a shift in bias, toward a greater propensity for reporting the absence of a signal, relative to the first pre-distractor block of trials (main effect of genotype on *b”_D_*: (F(2,24)=4.68, *P*=0.02, Cohen’s *d*=0.78; Fig. 7d). This genotype-specific response to the distractor remained stable throughout the remainder of the test session (no effect of block and no interaction between the effects of genotype and block; both F<1.80, both *P*>0.15). As was the case performing the unchallenged version of the task, all mice steadily omitted more trials in the course of the disruptor session (block 1 mean: 8.22%; block 5: 23.81%; main effect of block: F(1.82, 43.75)=11.59, *P*=0.0001; effects of genotype and interaction with block: both F<3.10, both *P*>0.06; not shown).

### Structural modeling of WT mouse CHT and the CHT Val89 variant

To gain insights into the structural underpinnings of the effect of the I89V substitution on the CHT transport mechanism, we pursued a template-based molecular modelling approach for WT mCHT and the Val89 variant. Our approach involved two key steps: identifying a suitable template and determining a high-quality sequence alignment between the template and query used during the production run. The use of hidden Markov model (HMM)-based profiles in fold-recognition algorithms, such as implemented in HHpred, allows for detecting distantly related homolog sequences. This approach identified the *Vibrio parahaemolyticus* sodium/sugar symporter (vSGLT, PDB id: 3DH4) (Faham et al., 2008) as the best template when compared with other SLC5 family members. The initial alignment of vSGLT and CHT covers 80% of the sequences with a sequence identity of 19%. Despite relatively low sequence similarity, the hydrophobicity profiles of the two transporters, used as indicators for proteins that share the same fold (Lolkema and Slotboom, 1998; Sarkar and Kellogg, 2010), showed visibly similar widths and distribution of hydrophobicity peaks, indicating that both proteins have a similar fold. It is noteworthy that TM14 in vSGLT has no counterpart in mouse (or human) CHT, which is predicted to have 13 transmembrane helices (Fig. 8a,b), extracellular N- and cytoplasmic C-termini (Fig. 8d).

**Figure 8.**
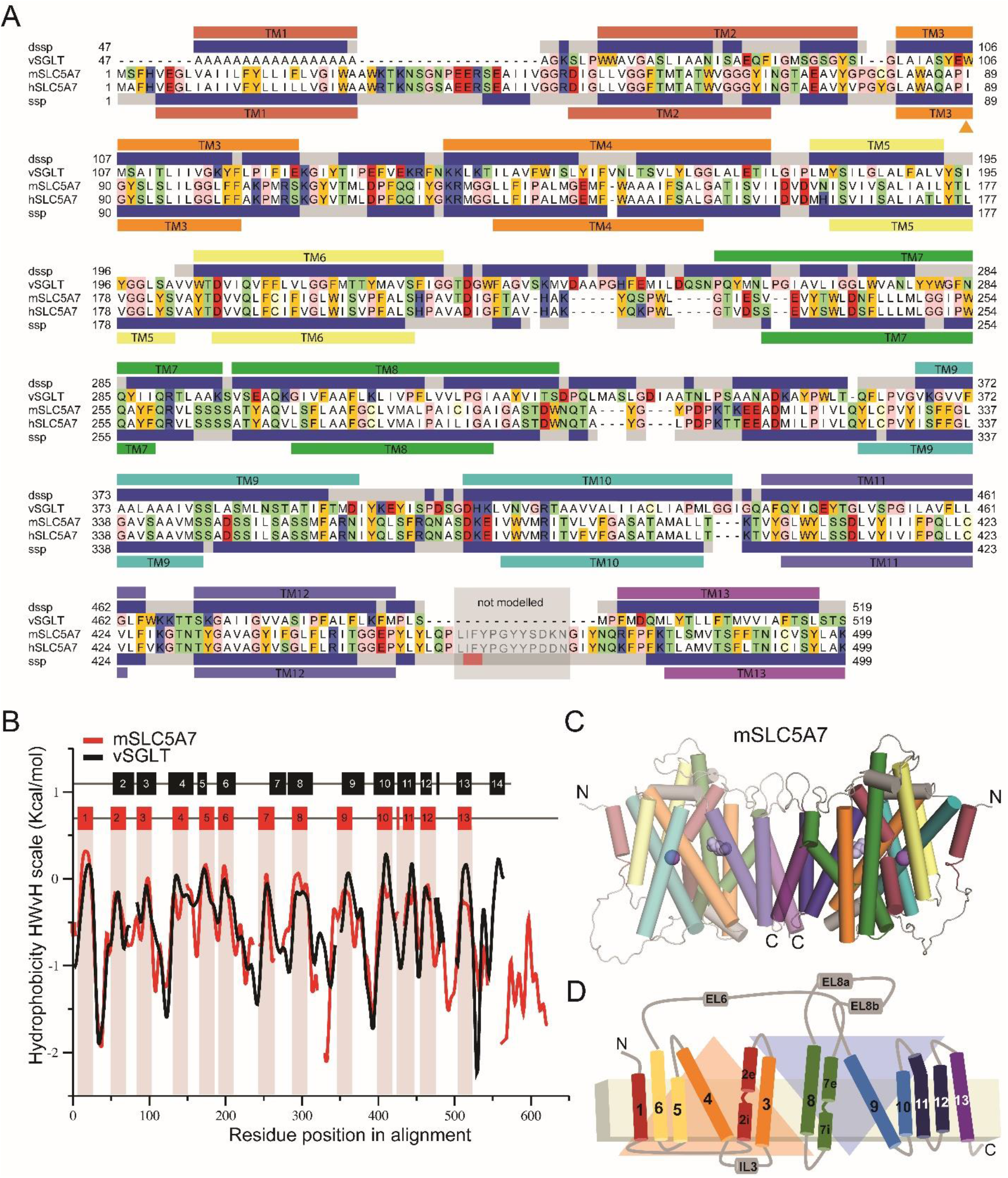
Structural model of a mouse CHT dimer. **a** Sequence alignment between mouse and human CHT (SLC5A7) and the template vSGLT used for structural modelling studies. The secondary structure prediction (SSP) averaged over the SLC5A7 profile from HHpred server (Hildebrand et al., 2009) is shown below hSLC5A7 sequence as gray (coil), blue (helix) and red (beta-strand) bars. The transmembrane topology prediction from Topcons server (Tsirigos et al., 2015) is also shown as bars below SSP colored and numbered according to the topology proposed by Faham et al. (2008). The secondary structure and transmembrane helices based on the structure of the template vSGLT are also shown on top of vSGLT sequence, similarly to that of CHT. The region between TM12 and TM13 shaded in gray was not modelled. Residues shown in the model are highlighted according to their properties; neutral in white, aromatic in yellow, polar in green, basic in blue and acidic in red. **b** Hydrophobicity profiles for mSLC5A7 (red) and vSGLT (black) revealing conserved hydrophobic regions. Residues corresponding to the TM helices in vSGLT or those predicted for mSLC5A7 are indicated as black and red bars at the top of the graph and in the case of mSLC5A7 also as red shading. **c,d** Final structural model of the dimeric form of mSLC5A7 and the correspondent topology. In both, TM helices are colored according to Faham et al. (2008) and the inverted repeat units are shown as orange and blue triangles.

The initial alignment was refined using an iterative procedure in which the gaps within secondary structural elements are removed, and the conservation scores from Consurf were used as a guide to position highly conserved residues within the core of the protein (Ashkenazy et al., 2016). The refined sequence alignment shows an overall correspondence between secondary structural elements, except in three of the loops connecting transmembrane helices. This matching was particularly striking for the first 13 TM helices (Fig. 8b). The broad sequence coverage between mCHT and vSGLT, similarity of the hydrophobicity profiles and matching between the TM helices, further substantiate the view that these two proteins share a common fold. The analysis of the stereochemistry for our model of CHT shows only one residue, located in a loop, in the disallowed region of the Ramachandran plot obtained with PROCHECK (see Methods). In addition to the excellent stereochemistry, the model has a ProQM score averaged by the number of residues (global ProQM score) of 0.67, comparable to that of the template (0.7), indicating the good quality of the model obtained.

Our work indicates that CHT has a similar global architecture as that of vSGLT (Fig. 8c,d). mCHT was modelled as a dimer, similar to the template, where transmembrane helices TM12 and TM13 constitute the dimerization interface with highly conserved residues taking part in the dimerization interacting network (Figs. 8c,d and 9a,b), supporting the possibility that mouse and human CHT assemble as a dimer. Each of the protomers has an inverted repeat topology, in which TMs 2-6 and TMs 7-11 constitute repeat units 1 and 2 (RU1, RU2) with a similar topology to the core of LeuT (TM1-10) (Faham et al., 2008), and a central cavity formed by TM2, 3, 7, 8 and 11 exposed to the cytoplasm, and helices 1, 12 and 13 as peripheral.

**Figure 9.**
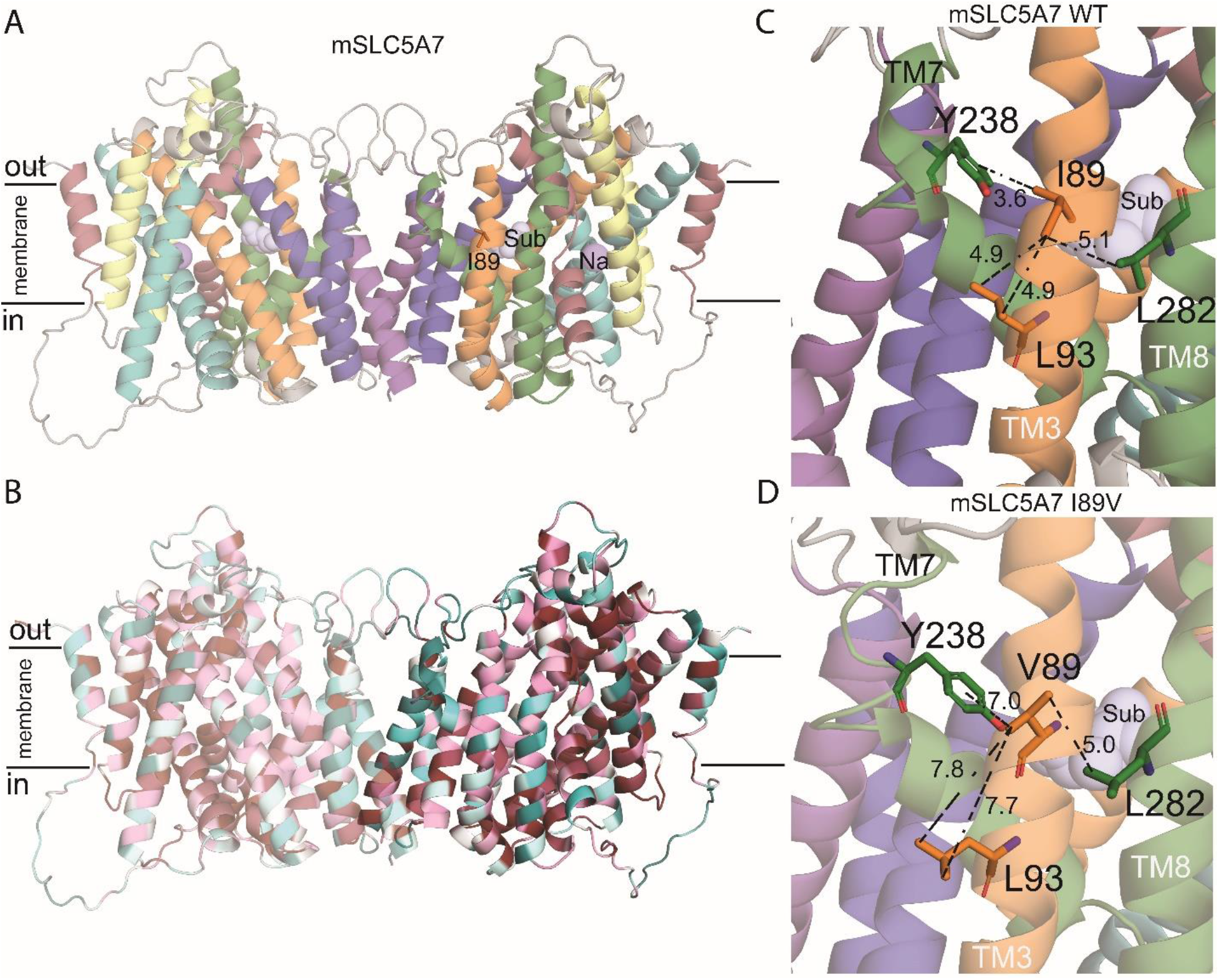
Structural model of mouse CHT Val89. **a** Structural model of mSLC5A7 in dimeric form colored according to the topology. I89, the sodium ion and substrate are highlighted as orange sticks and purple and light blue spheres, respectively. The position of the substrate and sodium ions where those obtained after structural superimposition of CHT model and vSGLT structure. **b** Final model of mSLC5A7 where each residue is colored by its conservation score obtained with Consurf server (Ashkenazy et al., 2016). **c,d** Close-up views of Ile89 and Val89 in their corresponding mSLC5A7 structures. Residues interacting with Ile89 are shown as sticks and the distances between sidechains are shown as dashed lines. The helices are colored as in **a**.

In our model, we found Ile89 to be located in TM3 of CHT, facing the membrane and stabilizing the packing between TM3, TM8 and TM7 by interacting with residues Leu93 (TM3), Tyr238 (TM7) and Leu282 (TM8) (Fig. 9c). This network of interactions is disrupted when Ile89 is substituted by Val89 (Fig. 9d), suggesting that the variant may destabilize helical packing in this region. It is worth noting that these three TMs are predicted to participate in shaping the cavity where choline binds, and thus Val89 may disrupt the exposure of the CHT substrate binding site. In addition, TM3, TM7 and TM8 are part of the “bundle” of the protein, which is believed to move with respect to the scaffold and the membrane plane upon the conformational changes responsible for transitions of the substrate binding site between states open to either extracellular or cytoplasmic spaces (alternating-access mechanism of transport). Destabilization of the fold next to position 89 by the Ile89Val substitution could also affect the ability of the transporter to adopt canonical conformations that the protein needs to reach during a transport cycle.

## Discussion

The current study aimed to determine the cholinergic impact of the common human CHT coding variant, Val89, that was previously demonstrated to be associated with an inattentive phenotype in humans, along with attenuated frontal activation during attentional performance (Sarter et al., 2016; Berry et al., 2015; Berry et al., 2014). The results indicate a dominant-negative reduction in choline clearance capacity in Val89 mice of approximately 80% in cortex and over 50% in striatum. Furthermore, although the initial depolarization-induced release of ACh from cholinergic terminals was similar across all mice, we found that 5 or 10 min following repeated depolarization, the capacity to release ACh was significantly attenuated in Val89 expressing mice. The effects of an attentional performance challenge indicated that, in contrast to WT mice, Val89 mice performed a continuous signal detection task largely through bottom-up, signal-driven processes. Immunoblotting indicated that surface expression of the CHT did not differ between the genotypes, suggesting that Val89-associated impairments were not due to aberrant surface trafficking. Structural modeling suggested potential cellular mechanisms underlying attenuated choline transport by the Val89 variant.

Results from the depolarization studies (Fig. 4) demonstrated that, in response to the initial depolarization, choline transport by the variant remains sufficient to support ACh release at a level that is comparable to WT mice. However, the impact of attenuated CHT function on ACh release capacity manifests following repeated depolarizations. Importantly, the paradigm we used to assess demonstrate a diminished the “reloading” capacity for subsequent ACh release involved a relatively small number of depolarizations at a relatively low frequency, mirroring ACh release events seen during hits in attentional task-performing rodents. An even greater impact of the Val89 variant would be expected when cholinergic neurons exhibit burst firing (e.g., Manns et al., 2003).

Our behavioral assessments indicated that in response to a visual disrupter, WT mice, but not Val89 mice, adopted a relatively more conservative criterion. Such a shift in bias is thought to reflect the deployment of a set of psychological processes that are conceptualized as voluntary top-down attentional control or model-based decision making (e.g., Clark et al., 2012). Specifically, in the present context, WT mice may have monitored disruptor-induced reward losses, triggering an increase in their threshold for reporting the presence of a signal. In contrast, Val89 mice may have relied primarily on the bottom-up detection of cues and model-free processing to perform this task. Consistent with the present evidence, such processing preserves unchallenged signal detection performance but limits the flexibility to respond to contextual alterations, such disruptor-induced performance loss.

The finding that WT mice adopted a more conservative bias in response to a disruptor was not associated with improvement in performance. This finding mirrors the absence of relatively greater performance deficits in heterozygous Val89 humans, compared to WT humans, using a comparable task and visual disruptor (Berry et al., 2015). We hypothesize that performing such a signal detection task may not sufficiently tax top-down control to produce performance loss arising from poor attentional control. To reveal performance benefits of a conservative criterion shift, and performance deficits in Val89 subjects, a rodent task that more severely taxes top-down processes and utilizes a true distractor, as opposed to a sensory disruptor, is needed, similar to the task used to demonstrate distractor vulnerability in Val89 humans (Berry et al., 2014).

The hypothesis that relatively poor attentional control results from attenuated cholinergic activity, is supported by evidence from rats with a spontaneous CHT intracellular trafficking deficit (Pitchers et al., 2017a; Pitchers et al., 2017b; Koshy Cherian et al., 2017; Sarter and Phillips, 2018; Phillips and Sarter, 2020; Paolone et al., 2013a). Furthermore, in patients with Parkinson’s Disease (PD), residual attentional performance, assessed by using a Continuous Temporal Expectancy Test (O’Connell et al., 2009) and challenged with a content-rich distractor, correlated with residual cholinergic innervation of cortex (Kim et al., 2019). Using the same test as in PD patients with cholinergic losses, humans heterozygous for the Val89 variant showed reduced resistance against such a distractor when compared with WT humans (Berry et al., 2014). In addition, Val89 humans encoded more distractor content than WT humans, further supporting relatively poor top-down attentional control as an essential phenotypic characteristic.

Our *in vivo* method indicated a relatively large reduction of CHT-mediated choline transport rates by the minor variant when compared with prior data obtained *in vitro* (Okuda et al., 2002). This finding warranted additional experiments to reveal the functionality of the CHT variant. The effects of HC-3 and lidocaine, while validating our *in vivo* choline transport measure in terms of a reduction in CHT-mediated transport, suggest either a residual, voltage-dependent choline transport capacity of CHT Val89 or a contribution to choline clearance by other electrogenic transporters (e.g., Onizuka et al., 2008). As interactions of choline, sodium, chloride and HC-3 with CHT Val89 have been reported to be unaffected in transfected cells (Okuda et al., 2002), our results are consistent with a diminished capacity for the high-affinity choline transport needed to sustain high-demand ACh release.

Our structural models of WT mouse CHT and the Val89 encoding transporter point to a structural importance of Ile89 in local packing of substrate transport-critical helices. We show that Ile89 is predicted to participate in an interacting network that stabilizes the fold of helices that contribute to the “bundle” domain of the “rocking bundle” model of transport (Forrest and Rudnick, 2009). This network may also contribute to the shaping of the binding site by interacting with TM7 and TM8. Because Ile89 is not predicted to contribute to the substrate binding site (see also Okuda et al., 2002), it seems most likely that any impact on choline recognition by Val89 must be indirect. Therefore, we favor a model whereby conformational transitions required to support alternating access of the choline binding site to extracellular and intracellular compartments are compromised. Additionally, our models were well fit to a dimeric structure solved for vSGLT, including TMs position to support the dimeric interface.

Such a structure may underlie dominant-negative mode of action, which has been previously shown for human disease-associated CHT coding variants (Barwick et al., 2012). Although choline clearance data from I/V mice showed compression toward zero, thereby limiting potentially the demonstration of greater effects in V/V mice, measures of potassium-evoked ACh release and response bias might have readily revealed gene-dose effects. In humans expressing Val89 variant, the presence of a gene-dose effect on attentional control remains unknown but appears unlikely given the already relatively large effect size seen in heterozygous Val89 humans (Fig. 2 in Berry et al., 2014). Further studies, including biophysical analyses of choline uptake and choline-induced currents (Iwamoto et al., 2006), are needed to characterize the conformational dynamics of the Val89 variant.

Here, we focused on measures of presynaptic cholinergic capacity in the cortex because of the association of the Val89 variant with heightened distractor vulnerability in humans and based on evidence indicating an essential role of cortical cholinergic activity for distractor filtering (references in Introduction). Although the demonstration of attenuated clearance rates in a second brain region, the striatum, is consistent with a systemic impact of the variant, the striatal alteration is less striking, indicating a region-dependence to the functional impact of Val89 that may also underlie why CHT Val89 mice show no overt behavioral, autonomic or motor deficits. Motor deficits would have drastically interfered with the ability to perform the basic operant response requirements of the signal detection task, including executing nose-pokes and retrieving water reward across relatively long test sessions and rapidly sequenced trials. We suspect that motor (and other peripheral) cholinergic terminals possess greater ACh synthesis reserve than those we queried in the CNS, though further efforts are needed to explore this possibility.

The present findings establish that the Val89 variant, which has previously been linked to attentional impairment in humans (Sarter et al., 2016; Berry et al., 2015; Berry et al., 2014; English et al., 2009), as well as major depression (Hahn et al., 2008), greatly impacts the capacity of cortical cholinergic presynaptic terminals to import choline and to sustain ACh release, even following relatively moderate depolarization rates. As such, the present evidence provides a mechanistic framework to interpret the increased distraction vulnerability and frontal cortical activation deficits in Val89-expressing humans. As impairments in cholinergically mediated, attentional-perceptual functions and frontal cortical abnormalities are essential variables in the manifestation of major psychiatric and neurological disorders (Kim et al., 2019; Berry et al., 2017; Lustig and Sarter, 2016; Albin et al., 2018; Sarter et al., 2021; Bohnen et al., 2009; Bohnen and Albin, 2009), and Val89 allele frequency is surprisingly common for a transporter functional coding variant, further investigation of the impact of the CHT Val89 variant in humans and CHT Val89 mice is warranted.

## Author contributions

ED and CA conducted the electrochemical experiments and analyzed the data. VP conducted immunoblotting experiments and analyzed the data. RDB generated the CHT Val89 mice and provided founders for the CHT Val89 breeding colony at the University of Michigan. SK trained and tested mice in the visual attention task and analyzed the data. ED, VP, RDB and MS designed the experiments. CFF developed the structural model of the CHT Val89 mutation. ED, CFF, RDB and MS wrote the original draft of the paper. All authors contributed to the editing of the final manuscript.

## Acknowledgement

We thank Evan Haley (Temple University) for assistance with subcellular fractionation and immunoblotting studies.

